# ComFB, a new widespread family of c-di-NMP receptor proteins

**DOI:** 10.1101/2024.11.10.622515

**Authors:** Sherihan Samir, Abdalla A. Elshereef, Vikram Alva, Jeanette Hahn, David Dubnau, Michael Y. Galperin, Khaled A. Selim

**Affiliations:** Interfaculty Institute of Microbiology and Infection Medicine, Organismic Interactions Department, Cluster of Excellence “Controlling Microbes to Fight Infections”, Eberhard Karls University of Tübingen, 72076 Tübingen, Germany; Department of Protein Evolution, Max Planck Institute for Biology Tübingen, Germany; Public Health Research Institute and Department of Microbiology, Biochemistry and Molecular Genetics, New Jersey Medical School, Rutgers University, New Jersey, USA; National Center for Biotechnology Information, National Library of Medicine, National Institutes of Health, Bethesda, Maryland 20894, USA; Institute of Phototroph Microbiology, Heinrich-Heine University Düsseldorf, 40225 Düsseldorf, Germany

## Abstract

Cyclic dimeric GMP (c-di-GMP) is a widespread bacterial second messenger that controls a variety of cellular functions, including protein and polysaccharide secretion, motility, cell division, cell development, and biofilm formation, and contributes to the virulence of some important bacterial pathogens. While the genes for diguanylate cyclases and c-di-GMP hydrolases (active or mutated) can be easily identified in microbial genomes, the list of c-di-GMP receptor domains is quite limited, and only two of them, PliZ and MshEN, are found across multiple bacterial phyla. Recently, a new c-di-GMP receptor protein, named CdgR or ComFB, has been identified in cyanobacteria and shown to regulate their cell size and, more recently, natural competence. Sequence and structural analysis indicated that CdgR is part of a widespread ComFB protein family, named after the “late competence development protein ComFB” from *Bacillus subtilis*. This prompted the suggestion that ComFB and ComFB-like proteins could also be c-di-GMP receptors. Indeed, we revealed that ComFB proteins from Gram-positive *B. subtilis* and *Thermoanaerobacter brockii* were able to bind c-di-GMP with high-affinity. The ability to bind c-di-GMP was also demonstrated for the ComFB proteins from clinically relevant Gram-negative bacteria *Vibrio cholerae* and *Treponema denticola*. These observations indicate that the ComFB family serves as yet another widespread family of bacterial c-di-GMP receptors. Incidentally, some ComFB proteins were also capable of c-di-AMP binding, identifying them as a unique family of c-di-NMP receptor proteins. The overexpression of *comFB* in *B. subtilis,* combined with an elevated concentration of c-di-GMP, suppressed motility, attesting to the biological relevance of ComFB as a c-di-GMP binding protein.

**IMPORTANCE:** The cellular content of the bacterial second messenger c-di-GMP is controlled by c-di-GMP synthases (GGDEF domains) and hydrolases (EAL or HD-GYP domains), whose activities, in turn, respond to the signals perceived by their upstream sensory domains. Cyclic-di-GMP transmits the signals to a variety of its targets, which may contain inactivated GGDEF, EAL, or HD-GYP domains, widespread PilZ or MshEN domains, or various lineage-specific c-di-GMP receptors. Many organisms encode multiple GGDEF domains but few c-di-GMP-binding proteins, suggesting the existence of still unidentified c-di-GMP receptors. Here, we demonstrate that the ComFB family proteins, which include the recently characterized cyanobacterial CdgR/ComFB, constitute yet another widespread family of bacterial c-di-NMP receptors. We additionally show that ComFB controls bacillar motility in a c-di-GMP-dependent manner.

## INTRODUCTION

Cyclic dinucleotide (c-di-NMP)-based second messengers are widely used by bacterial, archaeal, and eukaryotic cells to transduce signals from sensor proteins to various cellular receptors (Yoon & Waters, 2021). The first discovered c-di-NMP was 3’-5’-cyclic dimeric GMP (c-di-GMP), initially described in 1987 (Ross *et al*., 1987) and subsequently shown to be involved in protein and polysaccharide secretion, motility, cell division, cell development, and biofilm formation (Römling *et al*., 2013, Jenal *et al*., 2017), as well as contributing to the virulence of some important bacterial pathogens (Römling *et al*., 2013, Conner *et al*., 2017, Hall & Lee, 2018, Valentini & Filloux, 2019). Another second messenger, 3’-5’-cyclic dimeric AMP (c-di-AMP), was discovered in bacterial cells in 2008 (Witte *et al*., 2008, Corrigan & Gründling, 2013). Cyclic-di-AMP controls bacterial response to the changes in osmotic pressure and envelope stress by regulating the transport of potassium ions and glutamate, which makes it essential for cell growth in standard growth media (Commichau & Stülke, 2018, He *et al*., 2020, Krüger *et al*., 2021). In *Bacillus subtilis* and related bacteria, c-di-AMP also controls sporulation and is involved in the response to DNA damage (Commichau *et al*., 2015, Stülke & Krüger, 2020). In cyanobacteria, c-di-AMP primarily controls day-night metabolism via regulating glycogen synthesis, photosynthesis, redox balance and carbon/nitrogen metabolism (Selim *et al*., 2021a, Mantovani *et al*., 2022, Mantovani *et al*., 2023, Haffner *et al*., 2023b). Yet another second messenger, cyclic GMP-AMP (cGAMP), has been implicated in *Vibrio cholerae* virulence (Davies *et al*., 2012). In the past several years, cyclic dinucleotide second messengers have also been described in eukaryotes, where they are involved in cellular immunity and anti-viral defense (Millman *et al*., 2020, Yoon & Waters, 2021, Slavik & Kranzusch, 2023).

Cellular levels of c-di-GMP and c-di-AMP are controlled by environmental and intracellular signals that modulate the activities of the respective synthases (diguanylate and diadenylate cyclases, respectively) and hydrolases (phosphodiesterases). Genes encoding c-di-GMP synthetases (containing the GGDEF domain) and hydrolases (containing EAL or HD-GYP domains) are found in members of all bacterial phyla, sequenced so far (see the c-di-GMP census at https://www.ncbi.nlm.nih.gov/Complete_Genomes/c-di-GMP.html). In contrast to the well-conserved – and therefore easily identified – sequences of c-di-GMP turnover enzymes, c-di-GMP receptors are much harder to pinpoint (Chou & Galperin, 2016, Khan *et al*., 2023). So far, only two types of widespread c-di-GMP receptor proteins have been identified to contain PilZ or MshEN c-di-GMP-binding domains (Amikam & Galperin, 2006, Galperin & Chou, 2020, Wang *et al*., 2016, Junkermeier & Hengge, 2021, Sellner *et al*., 2021). The list of experimentally characterized c-di-GMP receptors also includes (i) c-di-GMP-binding riboswitches (Sudarsan *et al*., 2008); (ii) inactivated enzymes of c-di-GMP turnover, such as PelD, BcsE, or LapD, that contain modified (mutated or truncated) GGDEF, EAL, or HD-GYP domains (Whitney *et al*., 2012, Whitfield *et al*., 2020, Fang *et al*., 2014, Zouhir *et al*., 2020, Chatterjee *et al*., 2014), and (iii) a variety of lineage-specific proteins, such as BldD and RsiG (actinobacteria), FleQ (pseudomonads), VpsT and VpsR (vibrios), CLP-like transcriptional regulators (xanthomonads), or CheY-like response regulators with Arg-rich tails in *Caulobacter* sp. (Tschowri *et al*., 2014, Gallagher *et al*., 2020, Hickman & Harwood, 2008, Matsuyama *et al*., 2016, Waters *et al*., 2008, Krasteva *et al*., 2010, Chin *et al*., 2010, Nesper *et al*., 2017), reviewed in (Chou & Galperin, 2016, Khan *et al*., 2023). However, comparing the phylogenetic distribution of c-di-GMP turnover enzymes and c-di-GMP receptors reveals a variety of organisms whose genomes encode multiple GGDEF, EAL, and/or HD-GYP domains but very few, if any, c-di-GMP-binding proteins or riboswitches (see the above-mentioned c-di-GMP census web site). Further, the newly identified c-di-GMP-binding proteins typically have a narrow phylogenetic distribution (Skotnicka *et al*., 2020, Li *et al*., 2023, Nie *et al*., 2024). These discrepancies indicate the existence of previously unrecognized c-di-GMP receptors.

A generally similar picture is seen for c-di-AMP signaling. Genes encoding c-di-AMP-producing diadenylate cyclases are found in most bacteria and many archaea (Galperin, 2023), in keeping with the essentiality of this second messenger (Commichau & Stülke, 2018). Several c-di-AMP-binding domains have been described (Corrigan *et al*., 2013, He *et al*., 2020), e.g., various members of P_II_ signaling superfamily are conserved c-di-AMP-receptor proteins (Selim & Alva, 2024, Selim *et al*., 2023, Selim *et al*., 2021b, Selim *et al*., 2021a), but the current list of c-di-AMP-specific receptors is probably still incomplete (He *et al*., 2020, Stülke & Krüger, 2020, Mantovani *et al*., 2023).

Recently, a novel c-di-NMP receptor protein has been discovered in cyanobacteria and found to bind both c-di-GMP and c-di-AMP; it was initially named CdgR for “<u>c</u>-<u>d</u>i-<u>G</u>MP <u>R</u>eceptor” (Zeng *et al*., 2023). In the multicellular cyanobacterium *Nostoc* sp. PCC 7120 (Alr3277, GenBank accession number <u>BAB74976</u>), this 179-aa protein was shown to bind c-di-GMP with high affinity (Kd = 0.18 µM) and to regulate the bacterial cell size (Zeng *et al*., 2023). It could also bind c-di-AMP, albeit with a lower affinity (Kd = 2.66 μM). The CdgR homolog Slr1970 from the unicellular cyanobacterium *Synechocystis* sp. PCC 6803 showed weaker c-di-GMP binding (Kd = 1.68 µM) than Alr3277 but allowed solving the structure of its c-di-GMP-bound complex (Protein DataBank (PDB) entry <u>8HJA</u>). Slr1970 was found to bind also c-di-AMP with comparable affinity to c-di-GMP (Samir *et al*., 2023). We showed that both c-di-AMP and its receptor Slr1970 are required for DNA uptake and natural competence in *Synechocystis*. Since the CdgR proteins possess a close similarity to the late competence factor B domain from *Bacillus subtilis* (*Bs*ComFB; domain PF10719 in the Pfam database), we renamed Slr1970 to ComFB (also annotated as ComFB in the NCBI’s RefSeq database) (Samir *et al*., 2023). However, Zeng and colleagues argued that these proteins were functionally different (Zeng *et al*., 2023). In *B. subtilis*, the *comFB* gene is part of the *comF* competence operon that also encodes the DNA helicase ComFA and the predicted phosphoribosyltransferase ComFC, which mediate cellular uptake and handling of single-stranded DNA (Londono-Vallejo & Dubnau, 1993, Sysoeva *et al*., 2015, Damke *et al*., 2022). However, in contrast to these two neighbors, *Bs*ComFB appears to play no role in DNA uptake and its cellular role remains elusive (Sysoeva *et al*., 2015). Further, *comFB* genes are found in a variety of bacteria but missing in many *comFA*- and *comFC*-encoding organisms, including several *Bacillus* species (Kovacs *et al*., 2009, Kovacs *et al*., 2013). As a result, ComFB was assumed to participate in some kind of regulation. Given the widespread distribution of the ComFB-related proteins among diverse bacteria, we wondered if they could comprise a new family of c-di-GMP receptors that were missing in several bacterial lineages. Here, we present a bioinformatic and biophysical characterization of this protein family, along with binding assays of several of its members. This work shows that ComFB proteins, indeed, represent a widespread family of c-di-GMP (and also c-di-AMP) receptors. Moreover, we show its physiological relevance in controlling *B. subtilis* motility.

## RESULTS

### Phylogenetic distribution of ComFB proteins

To investigate the phylogenetic landscape of the ComFB family, we searched for homologs of the ComFB protein from *B. subtilis* (*Bs*ComFB) and CdgR from *Synechocystis* sp. PCC 6803 (Slr1970; *Sy*ComFB) using BLAST against the NCBI non-redundant protein sequence database. The search seeded with Slr1970 predominantly returned matches to other cyanobacterial proteins, while the search with *Bs*ComFB primarily yielded matches to proteins from the Bacillota phylum. Although these two proteins share the same core fold, their sequences are highly divergent, exhibiting less than 14% pairwise identity, with similarity largely limited to the residues involved in protein dimerization and the c-di-GMP-binding motif (Fig. 1A,B). Additionally, the zinc-binding cysteine residues found in *Bs*ComFB are not conserved in the cyanobacterial sequences (see below; Fig. 1B). To further substantiate the homology between *Bs*ComFB and *Sy*ComFB and to detect additional homologs, we ran HHpred searches against the PDB70 profile HMM database and the profile HMM database of various proteomes. Searches against representative proteomes identified ComFB family members in other phyla, including Actinomycetota, Bdellovibrionota, Myxococcota, Nitrospirota, Pseudomonadota, Spirochaetota, and Thermodesulfobacteriota (Table S1). HHpred also detected a match between *Bs*ComFB and *Sy*ComFB with a probability value greater than 99.5%, indicating their homology.

**Figure 1.**
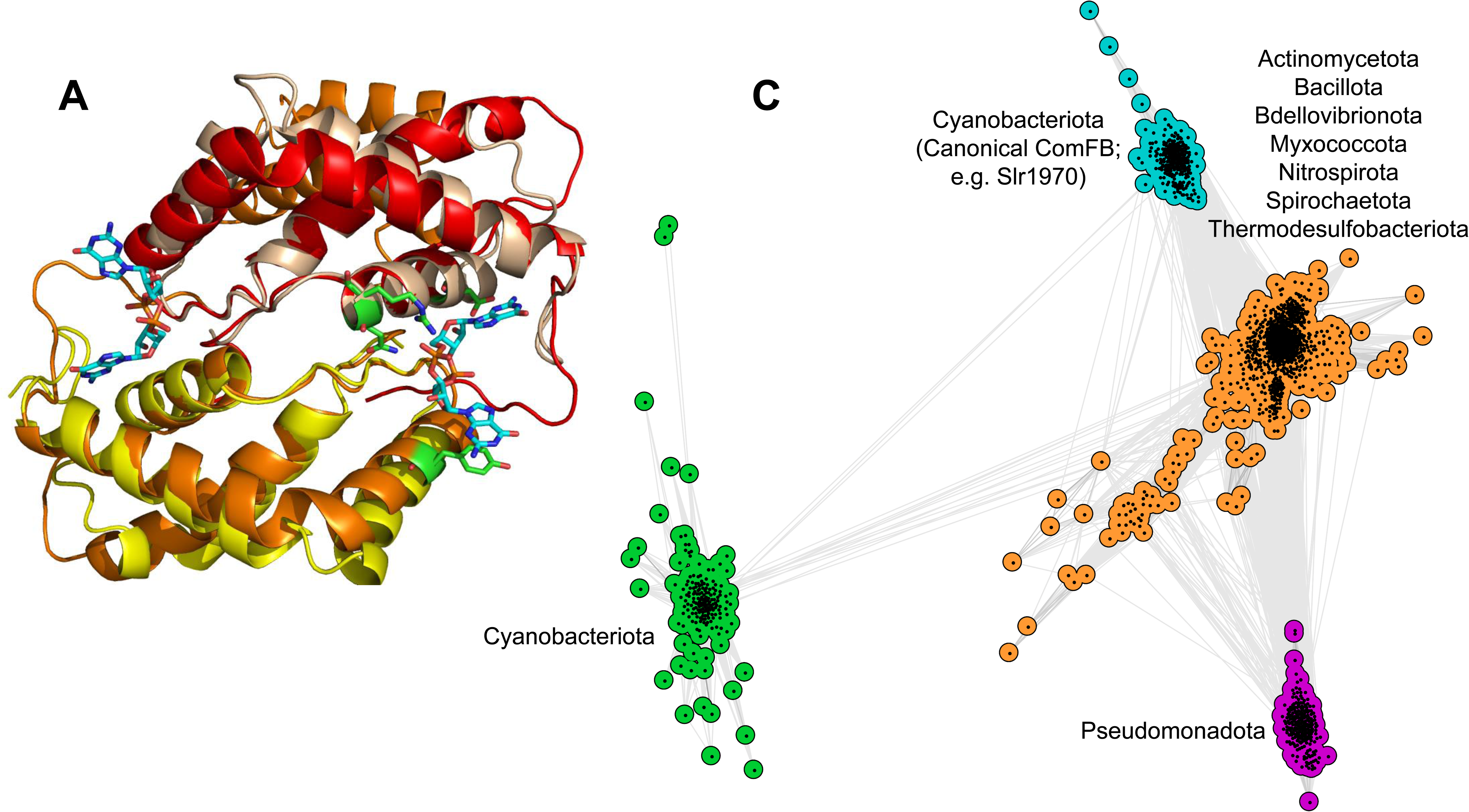
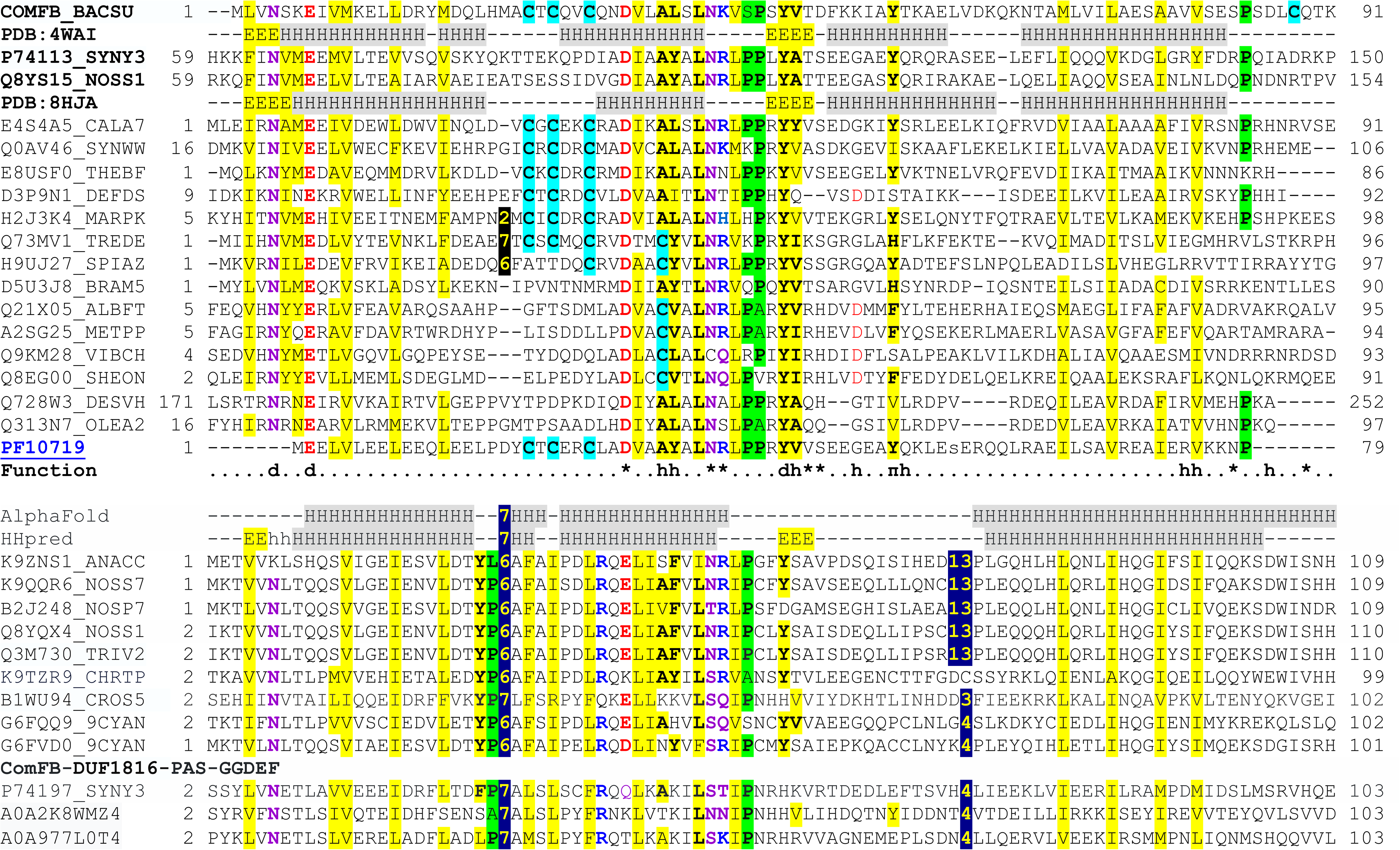
Sequence and structural conservation within the ComFB superfamily. (A) Structural alignment of the dimeric forms of *Bacillus subtilis* ComFB (*Bs*ComFB, PDB: <u>4WAI</u>, yellow and teal) and CdgR from *Synechocystis* sp. PCC 6803 (PDB: <u>8HJA</u>, orange and red). The CdgR-bound c-di-GMP molecules are shown in stick mode with carbon atoms in blue. The c-di-GMP binding residues D53, N100, R101 and Y115 of CdgR are shown in stick mode with carbon atoms in green. (B) Sequence alignment of representative members of the ComFB superfamily. Proteins are shown under their UniProt identifiers, and secondary structure assignments (H, α-helix, E, β-strand) of ComFB and CdgR are shown with their PDB codes. The numbers indicate the positions of the start and end of the alignment and the lengths of the gaps between the aligned blocks. Conserved negatively (D, E) and positively (K, R) charged residues are shown in red and blue, respectively; nonpolar hydrophilic residues (N, Q, S, T) are in purple. Conserved hydrophobic residues are indicated with yellow shading, and conserved turn residues (G, P, S, A) are shaded green. Zinc-binding Cys residues of ComFB and the conserved Cys residues in other proteins are shown on a light blue background. The last sequence in the upper block represents the Pfam entry PF10719. The symbols in the “Function” line indicate (as specified by Zeng *et al*., 2023): d, residues responsible for protein dimerization; asterisks, residues involved in binding c-di-GMP; h and π, residues involved in hydrophobic interactions with the c-di-GMP ligand. The lower block represents ComFB-related sequences that are not recognized by the PF10719 sequence model; its top two lines display secondary structure predictions by AlphaFold (Varadi *et al*., 2022) and HHpred (Zimmermann *et al*., 2018). The last three lines show sequences of the N-terminal ComFB domains of four-domain diguanylate cyclases. The sequences in the upper block are from the following organisms: ComFB, *Bacillus subtilis*; Q8YS15, *Nostoc* sp. PCC 7120; P74113, *Synechocystis* sp. PCC 6803 (both, cyanobacteria); E4S4A5, *Caldicellulosiruptor acetigenus*; Q0AV46, *Syntrophomonas wolfei*; E8USF0, *Thermoanaerobacter brockii* (all three, *Clostridia*); D3P9N1, *Deferribacter desulfuricans* (*Deferribacterota*); H2J3K4, *Marinitoga piezophila* (*Thermotogota*); Q73MV1, *Treponema denticola*; H9UJ27, *Spirochaeta africana*; D5U3J8, *Brachyspira murdochii* (all three, *Spirochaetota*); Q21X05, *Albidiferax ferrireducens*; A2SG25, *Methylibium petroleiphilum* (both, *Betaproteobacteria*); Q9KM28, *Vibrio cholerae*; Q8EG00, *Shewanella oneidensis* (both, *Gammaproteobacteria*); Q728W3, *Desulfovibrio vulgaris*; Q313N7, *Oleidesulfovibrio alaskensis* (both, *Thermodesulfobacteriota*). All sequences in the lower block are from cyanobacteria. (C) Cluster map of ComFB homologs. A set of 1,626 representative ComFB sequences (≤ 70% pairwise identity and ≥ 70% length coverage) was clustered using the CLANS tool (Frickey & Lupas, 2004) based on pairwise BLAST P-values. Dots represent individual sequences, colored according to their group. Line color intensity reflects sequence similarity, with darker lines indicating higher similarity. The analysis revealed four clusters: two within Cyanobacteriota, one comprising diverse phyla (e.g., Actinomycetota, Bacillota), and a distinct Pseudomonadota cluster, highlighting conserved c-di-GMP-binding residues across these diverse groups.

The structures of *Bs*ComFB (PDB: <u>4WAI</u>) and Slr1970 (PDB: <u>8HJA</u>) represent stable dimers, with each subunit consisting of four long α-helices and one or two short β-strands (Fig. 1A,B). Additionally, Slr1970 features an N-terminal three-helical subdomain, which is almost exclusively found in its cyanobacterial homologs, as well as in some uncharacterized cyanobacterial proteins, such as *Nostoc punctiforme* Npun_F5121 (UniProt: B2J269). The structures of *Bs*ComFB and Slr1970 align with an RMSD of 3.6 Å over 91 C_α_ residues, despite their low (14%) pairwise sequence identity.

Using structure-based sequence alignment of *Bs*ComFB and Slr1970 in iterative database searches, we identified members of this protein family in several distinct bacterial lineages. These lineages include, in addition to cyanobacteria, members of the phyla Bacillota (classes Bacilli, Clostridia, Negativicutes, and Tissierellia), Pseudomonadota, Deferribacteriota, Spirochaetota, Thermotogota (Fig. 1B), and several others, as well as various candidate phyla (see a larger alignment in Fig. S1 and a partial list of family members in Table S1). A representative selection of ComFB family proteins is also available in the InterPro database (Paysan-Lafosse *et al*., 2023) under entry <u>IPR019657</u>, corresponding to the Pfam database (Mistry *et al*., 2021) domain PF10719. However, we discovered a subfamily of ComFB-related proteins widespread in cyanobacteria (Fig. 1C), but not included in the InterPro database, likely due to the presence of two insertions and changes in several conserved residues, including the c-di-GMP binding motif (Fig. S1 and Table S1). Most members of this new subfamily, such as, *Nostoc* sp. All3687 (UniProt: Q8YQX4), *Anabaena cylindrica* Anacy_5104 (UniProt: K9ZNS1), *Trichormus variabilis* Ava_3600 (UniProt: Q3M730) and CEN44_21400 of *Fischerella muscicola* (UniProt: A0A2N6JYB2, misannotated as a DUF3349 domain-containing protein), represent stand-alone ComFB-like domains. However, in several proteins within this new subfamily, such as Sll1170 from *Synechocystis* sp. (UniProt: P74197), the N-terminal ComFB-like domain is followed by DUF1816, PAS, and GGDEF domains. This domain architecture suggests potential diguanylate cyclase activity, with the ComFB-like domain presumably acting as a regulatory domain.

Hereafter, we refer to these proteins collectively as the ComFB family, as the name CdgR is potentially confusing, having been previously used for the c-di-GMP regulator YdiV, which consists of a stand-alone EAL domain (Hisert *et al*., 2005, El Mouali *et al*., 2017).

Most bacterial genomes encode a single copy of the *comFB* gene, if any (Table S1). However, certain lineages, particularly within Clostridia, Cyanobacteria, and Spirochaetota, often encode two or more orthologs of ComFB. Specifically, the cyanobacterial genomes of *Synechococcus* sp. PCC 7502, *Pseudanabaena biceps* PCC 7429, and *Pseudanabaena* sp. PCC 7367 encode six ComFB proteins each (Table S1). *Synechocystis* sp. PCC 6803 possesses four ComFB family members (Slr1970, Slr1505, Sll1739, and Sll1170), while *F. muscicola* CCMEE 5323 and *Nostoc* sp. PCC 7120 each has two different ComFB-like proteins.

Outside bacterial lineages, ComFB family members have been found in two archaea, *Methanocella* sp. (GenBank: <u>OPY29888</u>) and *Candidatus* Methanomethylicus sp. (GenBank: <u>HGS80389</u>, recently suppressed but still available in UniProt and on the NCBI and EBI websites), the fornicate *Aduncisulcus paluster* (GenBank: <u>GKT31366</u>), and as part of a multidomain protein from the model plant *Arabidopsis thaliana* (GenBank: <u>OAO89096</u>; see Fig. S1). These instances likely arose from bacterial contamination.

In many lineages, ComFB exhibits a patchy distribution, including its previously noted presence in close relatives of *B. subtilis* but not in other *Bacillus* species such as *B. anthracis, B. cereus*, and *B. thuringiensis* (Kovacs *et al*., 2009, Kovacs *et al*., 2013). Within the phylum *Pseudomonadota*, ComFB is found in the classes Betaproteobacteria and Gammaproteobacteria (Fig. S1), as well as in the former Deltaproteobacteria (recently reclassified as the phylum Thermodesulfobacteriota), but so far not in Alphaproteobacteria or Epsilonproteobacteria (the new phylum Campylobacterota). Furthermore, all betaproteobacterial ComFB sequences come from the single order Burkholderiales, and nearly all deltaproteobacterial sequences are from the order Desulfovibrionales (Table S1). Among gammaproteobacteria, ComFB sequences are found in lineages such as Aeromonadales, Alteromonadales, Pseudomonadales, and Vibrionales, but so far not in enterobacteria or xanthomonads. This patchiness is also evident at lower taxonomic levels. For example, within the order Pseudomonadales, all ComFB sequences are found in the genus *Marinobacter* and none in *Pseudomonas*. Similarly, among spirochetes, ComFB is encoded in oral pathogens *Treponema denticola* (Fig. 1B) and *Treponema socranskii*, as well as in the cattle skin pathogen *Treponema brennaborense*, but not in the closely related *Treponema pallidum*, the causative organism of syphilis, or the Lyme disease spirochete *Borrelia* (*Borreliella*) *burgdorferi* (Table S1).

### Evolutionary landscape of ComFB superfamily

To gain a comprehensive understanding of the evolutionary relationships among ComFB family proteins, we gathered homologs of several representative ComFB proteins and clustered them using the CLANS tool (Frickey & Lupas, 2004) based on the strength of their all-against-all pairwise sequence similarities. The resulting map (Fig. 1C) revealed four distinct clusters of ComFB sequences: two exclusively composed of proteins from the Cyanobacteriota phylum, one from the Pseudomonadota phylum, and one that includes proteins from a variety of phyla. Despite the low sequence similarity (less than 15%) between these clusters, many residues of the c-di-GMP-binding motif are conserved (Fig. 1B).

The *B. subtilis* ComFB sequence is located in the central cluster (colored orange), which includes sequences from diverse phyla such as Actinomycetota, Bacillota, Bdellovibrionota, Myxococcota, Nitrospirota, Spirochaetota, and Thermodesulfobacteriota, and from which the other three clusters are radiating. The *Synechocystis* ComFB sequence (Slr1970; *Sy*ComFB), along with its additional homologs in *Synechocystis* (Slr1505 and Sll1739), are in the closely connected cyanobacterial cluster (in cyan). This cluster also contains the *Nostoc* CdgR/ComFB (Alr3277), the multiple ComFB homologs found in *Synechococcus* sp. PCC 7502, *Pseudanabaena* biceps PCC 7429, and *Pseudanabaena* sp. PCC 736, as well as the fusions of the ComFB-like domain with phage shock protein A (*Synechococcus* sp. PCC 7502; UniProt: K9SSQ8) and ABC transporter ATP-binding protein (*Limnothrix* sp. P13C2; UniProt: A0A1C0VII2).

The second cyanobacterial cluster (in green), which is weakly connected to the central cluster (in orange), contains the new subfamily of ComFB-like proteins (lower sequence block in Fig. 1B). This cluster consists of stand-alone ComFB-like proteins (e.g., CEN44_21400, All3687, and Ava_3600) and multidomain diguanylate cyclases with an N-terminal ComFB-like domain, such as the *Synechocystis* diguanylate cyclase Sll1170 (UniProt: P74197). The separation of this new cyanobacterial subfamily in the map suggests functional distinction from the other cyanobacterial ComFB proteins. Notably, many cyanobacterial species have representatives in both cyan and green clusters (e.g., *Synechocystis* sp. PCC 6803, *Nostoc* sp. PCC 7120, *F. muscicola,* and *T. variabilis* ATCC 29413). The fourth cluster (in violet) possesses sequences of Pseudomonadota phylum, and most sequences of this cluster consist solely of the ComFB domain.

### Sequence conservation within the ComFB family

Despite the relatively low level of sequence identity between *Bs*ComFB and *Sy*ComFB, the key functional residues are conserved between these two (Fig. 1B) and within the entire family (Fig. S1). Indeed, the three residues, Asn4, Glu7, and Tyr46 (*Bs*ComFB numbering), that are involved in protein dimerization, are conserved in ComFB proteins from all bacterial phyla (Fig. S1 and Fig. S2). Accordingly, it would be reasonable to assume, that, like *Bs*ComFB and *Sy*ComFB (Zeng *et al*., 2023, Sysoeva *et al*., 2015), all family members form dimers (or higher oligomers). In addition, several c-di-GMP-binding residues, identified in the structure of the *Sy*ComFB-c-di-GMP complex (PDB: 8HJA) (Zeng *et al*., 2023), are widely conserved as well (Fig. 1B and Fig. S1). These residues include (i) Asp33, Asn40, and Arg/Lys41 (*Bs*ComFB numbering) that bind c-di-GMP through electrostatic interactions; (ii) hydrophobic amino acid residues in positions of Ala36, Leu37, and Val47 that form hydrophobic contacts with the c-di-GMP ligand, and (iii) Tyr55 that forms a π-bond with the guanine moiety of c-di-GMP. Several other c-di-GMP-binding residues of *Sy*ComFB are poorly conserved, suggesting greater flexibility in ligand binding within the family with altered affinity (or a loss of binding altogether).

### *comFB* genomic neighborhoods

Members of the ComFB family are typically annotated as “late competence development protein ComFB”, based on *Bs*ComFB, the founding member of the family (Londono-Vallejo & Dubnau, 1993, Sysoeva *et al*., 2015). However, the *comFA*-*comFB-comFC* operon organization is only seen in *B. subtilis* (Fig. 2) and its closest relatives, *B. amyloliquefaciens*, *B. atrophaeus*, *B. licheniformis*, *B. pumilis, B. velezensis*, and *B. xiamenensis*. A comparison of the *comF* gene distribution using the COG database (Galperin *et al*., 2021) shows that *comFB* is typically found without either *comFA* or *comFC*, and often in those organisms that do not carry either of these genes (Table S2).

**Figure 2.**
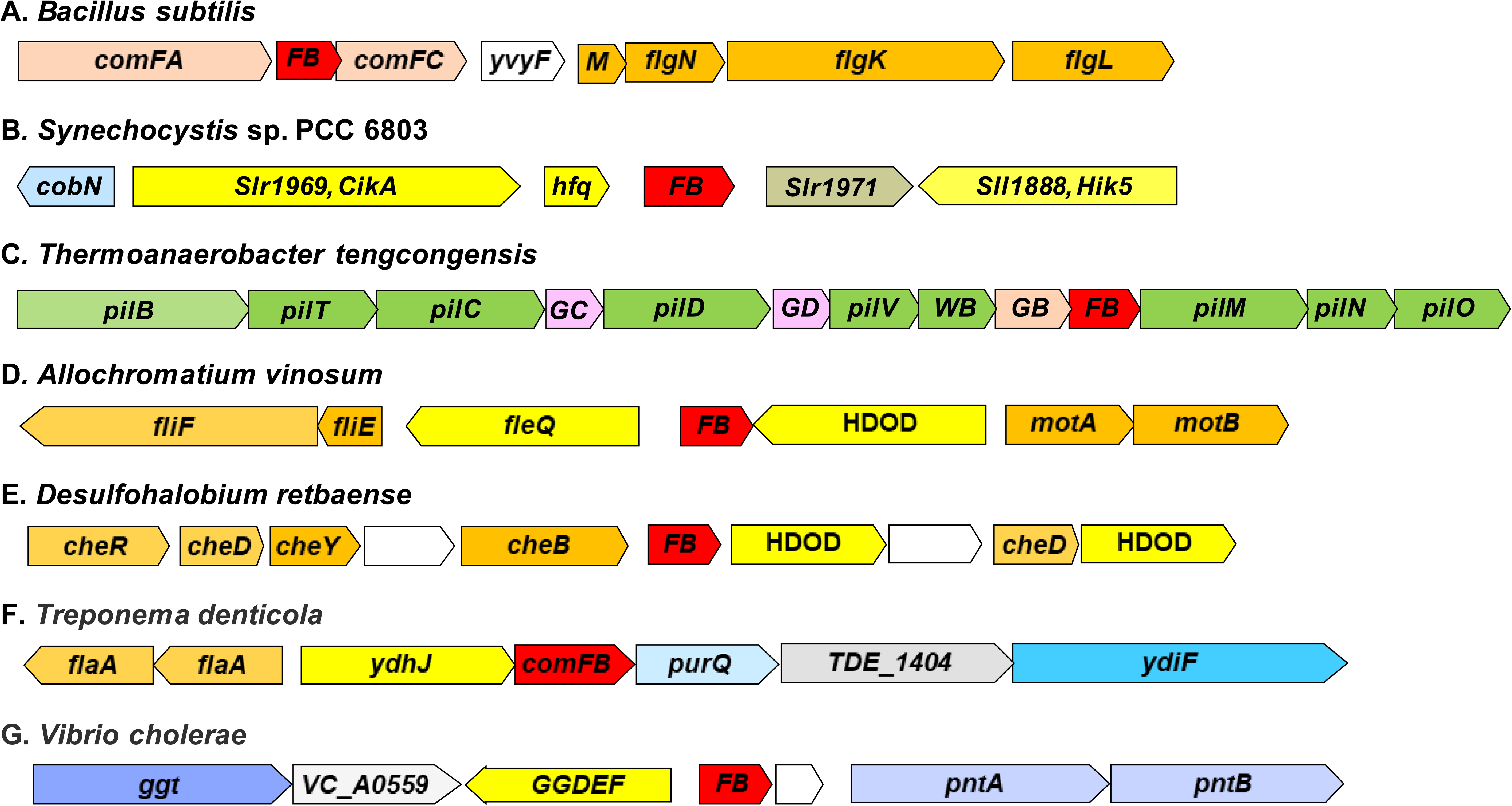
Genomic neighborhoods of selected ComFB/CdgR family proteins. Genomic fragments are listed with the organism names, GenBank accession numbers, and genomic coordinates. Gene sizes are drawn approximately to scale, and gene names are from GenBank, RefSeq, and/or the COG database. *ComFB* genes are in red, other competence-related genes are in pink, flagella-related genes are in orange, pili-related genes are in green, signal transduction genes are in yellow, metabolic genes are in various shades of blue, poorly characterized genes are in grey or white. The graph displays fragments of the following genomes: A, *Bacillus subtilis* 168, GenBank accession AL009126: 3,643,558..3,637,338; B, *Synehocystis* sp. PCC 6803, BA000022: 1,776,983..1,783,355; C, *Thermoanaerobacter tengcongensis* MB4, AE008691: 1,261,742..1,271,495; D, *Allochromatium vinosum* DSM 180, CP001896: 3,319,557..3,327,133; E, *Desulfohalobium retbaense* DSM 5692, CP001734: 792,659..799,553; F, *Treponema denticola* ATCC 35405, AE017226: 1,452,441..1,444,590; G, *Vibrio cholerae* O1 biovar El Tor str. N16961 chromosome II, AE003853: 495,055..503,153. The genomic fragments for *B. subtilis* and *T. denticola* are shown in reverse complement.

In *B. subtilis* and several other organisms (e.g. *Treponema denticola* ATCC 35405), *comFB* is located in the vicinity of one or more flagellar genes (Fig. 2). In many members of the clostridial order Thermoanaerobacterales, including *Caldicellulosiruptor* and *Thermoanaerobacterium* species, *comFB* is located within the operon that codes for type IV pili (Khan *et al*., 2020) (see Fig. 2). In cyanobacteria, *comFB* is frequently found in an operon with *hfq* (Fig. 2), a key component of cyanobacterial type IV-pili machinery that is required for motility and DNA uptake (Samir *et al*., 2023, Schuergers *et al*., 2014). This genomic organization suggests a possible involvement of ComFB proteins in the regulation of cell motility or other type IV pili-related functions. In *Vibrio cholerae*, *V. harveyi*, *V. parahaemolyticus* and some other organisms (e.g. *Aliivibrio fischeri*), the *comFB* gene is transcribed divergently from a gene coding for a GGDEF-containing diguanylate cyclase, highlighting a possible functional link to c-di-GMP signaling.

### ComFB domain architectures

The vast majority of ComFB domains are found in a stand-alone form (e.g., *Bs*ComFB; PDB: 4WAI) (Fig. 3). Also, most sequences of the Pseudomonadota cluster, e.g. as seen in *V. cholerae*, are found as a sole ComFB domain (see the violet cluster in Fig. 1C). However, canonical cyanobacterial members often contain an additional three-helical N-terminal subdomain, as seen in the structure of *Sy*ComFB (PDB: 8HJA), or other N-terminal and/or C-terminal extensions, as in *Synechocystis* sp. PCC 6803 proteins Slr1505 and Sll1739 (see Fig. 3 and Fig. S3 for domain architectures and GenBank accession numbers). Some cyanobacterial proteins contain two tandem ComFB domains (e.g., Pse7367_0880, UniProt: K9SGA4) (Fig. 3). Remarkably, such tandem-domain ComFB proteins include five out of the six paralogs encoded by the genomes of *Pseudanabaena biceps* PCC 7429, *Pseudanabaena* sp. PCC 7367, and *Synechococcus* sp. PCC 7502 (Table S1 and Fig. S3). Some of these proteins (such as UniProt: K9SSQ8) additionally contain long N-terminal coiled-coil segments that bear limited similarity to phage shock protein A of the PspA/IM30 family (Pfam domain PF04012) (Fig. 3 and Fig. S3).

**Figure 3.**
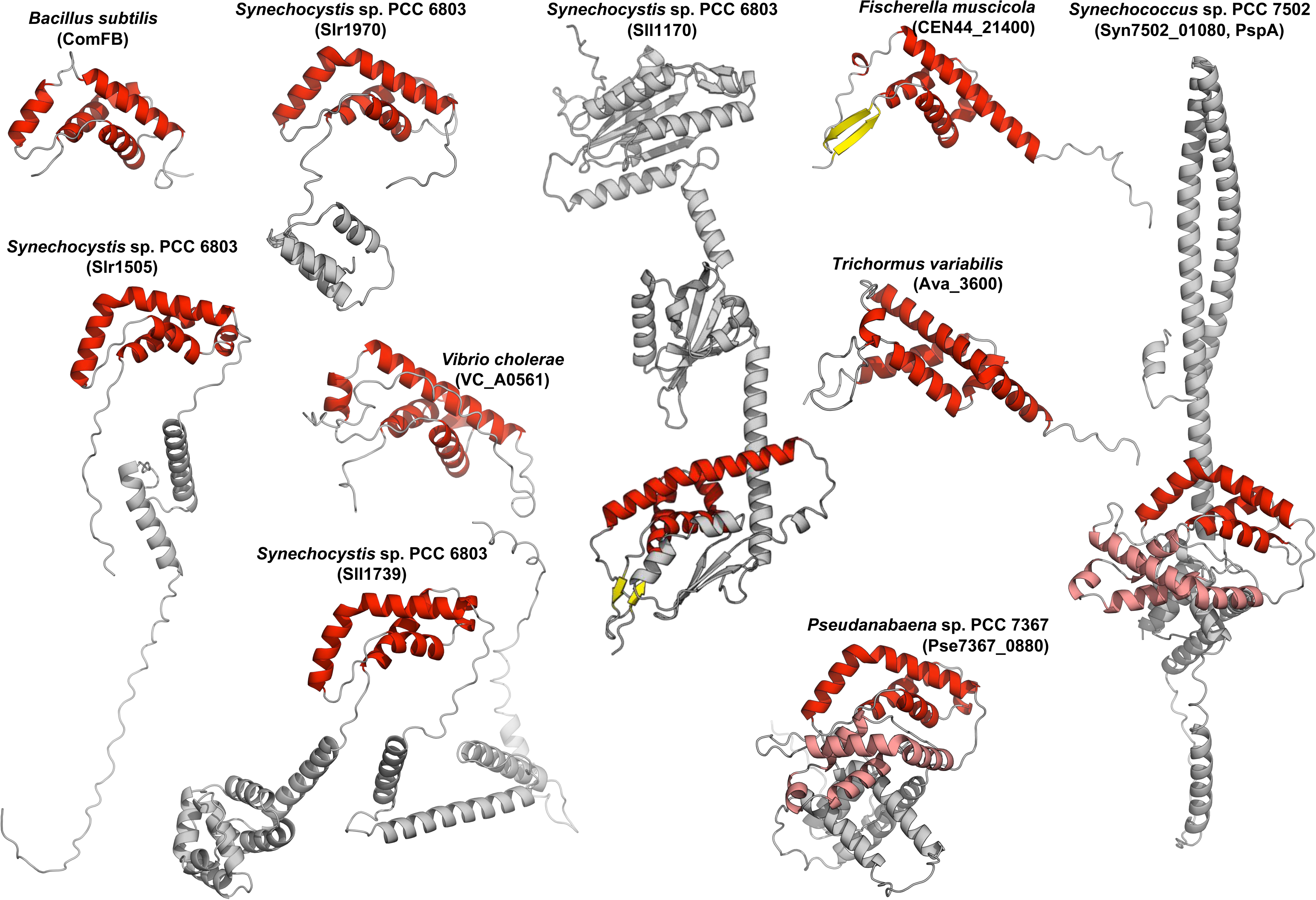
Structural gallery of representative ComFB domain-containing proteins from various species. α-helices in the ComFB domain are colored red, β-strands are in yellow, and the remainder of the protein in grey. For proteins with two ComFB domains, one domain is shown in lighter shades. The structures are AlphaFold2 (Jumper *et al*., 2021) predictions from UniProt/AlphaFold DB (Varadi *et al*., 2024), except for *Bacillus subtilis* (PDB 4WAI). The species represented include *Synechocystis sp.* PCC 6803 (Slr1970, Sll1170, Slr1505, and Sll1739, UniProt accessions P74113, P74197, P73943, and P73385, respectively), *Vibrio cholerae* (Q9KM28), *Fischerella muscicola* (A0A2N6JYB2), *Trichormus variabilis* (Q3M730), *Synechococcus sp.* PCC 7502 (K9SSQ8), and *Pseudanabaena sp.* PCC 7367 (K9SGA4). Several of these proteins also contain additional features, such as N- or C-terminal extensions (e.g., Slr1505), coiled-coil segments (e.g., PspA), or other domains: Sll1739 and Slr1970 have uncharacterized α-helical bundle domains, and Sll1170 contains DUF1816, PAS, and GGDEF domains.

Several other cyanobacteria, such as *Halothece* sp. PCC 7418, combine ComFB (e.g., UniProt: K9YBB1) with an N-terminal PATAN (DUF4388, <u>PF14332</u>) domain (Fig. S4), which is usually found in PatA-type response regulators that control heterocyst formation in cyanobacteria (Makarova *et al*., 2006). Remarkably, a large-scale protein interaction screen in *Synechocystis* sp. PCC 6803 revealed an interaction between the second ComFB protein, encoded by *slr1970*, and the Slr1594 protein of the PatA family (Sato *et al*., 2007); however, the relevance of this interaction is still unclear. Additionally, in the cyanobacterium *Limnothrix* sp. P13C2, the ComFB domain is found in combination with an ATPase subunit of an ABC transporter (UniProt: A0A1C0VII2) (Fig. S3). Also, as mentioned above, in the Sll1170 protein of the cyanobacterium *Synechocystis* sp. PCC 6803 and several other proteins, the N-terminal ComFB domain is followed by DUF1816, PAS, and GGDEF domains (Fig. 3 and Fig. S3).

Other architectures can also be found (Fig. S3), including several clostridial response regulators that combine the ComFB domain and the two-component phosphoacceptor receiver (REC, <u>PF00072</u>) domain (e.g. UniProt: R6WME1) (Fig. S3 and Fig. S4). Similarly, some proteins from the *Candidatus* Omnitrophota phylum (e.g., UniProt: A0A1G1PT98) couple the ComFB domain with the chemotaxis methyltransferase CheR (<u>PF01739</u>) domain (Fig. S3 and Fig. S4). The ComFB proteins from *Treponema denticola* (UniProt: Q73MV1) and many other spirochetes combine an N-terminal ComFB domain with a predicted C-terminal immunoglobulin-like (transthyretin-like) domain (Fig. S3 and Fig. S4; Table S1). This diverse array of domain architectures highlights the functional versatility of the ComFB family and suggests that these proteins may play a variety of regulatory roles across different bacterial lineages.

### Metal-binding Cys residues in the ComFB family

In the structure of ComFB from *B. subtilis*, each monomer contains a tightly bound Zn^2+^ ion that is coordinated by four Cys residues: Cys25, Cys27, Cys30, and Cys88, and appears to contribute to the stabilization of the protein (Sysoeva *et al*., 2015). Examination of ComFB sequences from other members of the phylum *Bacillota* reveals conservation of the first three Cys residues but not the last one (Fig. 1B). These three Cys residues are also conserved in ComFB proteins from members of the phyla Deferribacteres and Thermotogae and occasionally found in proteins from other lineages, such as *Maridesulfovibrio salexigens* (Thermodesulfobacteriota) and the cyanobacterium *Synechococcus* sp. JA-2-3B’a(2-13) (Fig. S1). The same three Cys residues are conserved in some, albeit not all, spirochetes. Some spirochete ComFB sequences, like the one from *T. denticola*, contain an additional Cys residue that is located in the last turn of the second long α-helix. Others, like ComFB from *Spirochaeta africana*, carry just two of these Cys residues, and some, like ComFB from *Brachyspira murdochii*, neither of them (Fig. 1B). Beta- and gammaproteobacterial ComFB sequences typically contain a single conserved Cys residue in the same position, near the end of the second long α-helix. Finally, except for the above-mentioned *M. salexigens* and *Synechococcus* sp. JA-2-3B’a(2-13), ComFB proteins from the phyla Cyanobacteriota and Thermodesulfobacteriota do not contain any of these Cys residues. Altogether, the conservation of these Cys residues is seen only in a fraction of ComFB family members (Fig. S2).

### Cyclic-di-NMP binding by ComFB proteins

Although c-di-GMP binding by *Sy*ComFB (Slr1970) was suggested to be specific for cyanobacteria (Zeng *et al*., 2023), the structural similarity between *Sy*ComFB and *Bs*ComFB (Fig. 1A), coupled with the conservation of several c-di-GMP-binding residues within the entire ComFB family (Fig. 1B), suggested that other members of this family might also serve as c-di-GMP and/or c-di-AMP receptors. Moreover, we were able to show that SyComFB can also bind c-di-AMP with comparable affinity to c-di-GMP (Samir *et al*., 2023). To investigate whether c-di-NMP binding is a common property of diverse members of the ComFB family, we heterologously expressed ComFB proteins from four phylogenetically distinct bacterial lineages and tested their ability to bind c-di-GMP and/or c-di-AMP. The binding assays were performed with ComFB proteins from *B. subtilis* (phylum Bacillota, class Bacilli), *Thermoanaerobacter brockii* (phylum Bacillota, class Clostridia), *Vibrio cholerae* (phylum Pseudomonadota, class Gammaproteobacteria), and *Treponema denticola* (phylum Spirochaetota, class Spirochaetia); their sequences are shown in Fig. 1B. While both *B. subtilis* and *T. brockii* are environmental Gram-positive bacteria, *V. cholerae* and *T. denticola* represent pathogenic Gram-negative bacteria.

First, we used size exclusion chromatography coupled to multiangle light scattering (SEC-MALS) to determine the oligomeric state of all ComFB proteins in solution. As expected, *B. subtilis* ComFB (*Bs*ComFB) protein showed a species of ∼ 23.5 kDa (Fig. S5), indicating that *Bs*ComFB protein (the theoretical molecular mass of a monomer with an 8xHis tag is 12.1 kDa) is a dimer in solution, in agreement with the crystal structure of *Bs*ComFB (Sysoeva *et al*., 2015). Similarly, *T. denticola* ComFB (*Td*ComFB; with the theoretical mass of monomer of 27.8 kDa) behaved as a dimer in solution with a molar mass of 45.5 kDa (Fig. S5). Unexpectedly, ComFB proteins of both *T. brockii* (*Tb*ComFB) and *V. cholerae* (*Vc*ComFB) behaved as monomers in solution with molar masses of 11.4 kDa and 13.4 kDa, respectively (Fig. S5).

Next, we measured the binding affinity of ComFB proteins to either c-di-GMP or c-di-AMP using isothermal titration calorimetry (ITC). The raw isothermal data for titration of c-di-GMP or c-di-AMP were fitted using a one-binding site model for monomeric ComFB proteins to determine the dissociation constant (K_D_) (Table 1; Fig. 4 and Fig. 5). For *Bs*ComFB, the titration of c-di-GMP yielded an exothermic profile with a high affinity in the very low micromolar range (K_D_ 0.17 μM; Fig. 4A). In contrast, the c-di-AMP binding events showed endothermic calorimetric signals with a K_D_ of 83.3 μM, indicating very weak binding (Fig. 5), implying that *Bs*ComFB preferentially binds to c-di-GMP. Next, we assessed the ability of *Bs*ComFB protein to bind to c-di-GMP in the presence of saturating concentrations of c-di-AMP, and *vice versa*. In a competition binding assay, when *Bs*ComFB protein was saturated first with 150 µM c-di-AMP, c-di-GMP binding events again showed strong exothermic calorimetric signals, however the binding enthalpy of c-di-GMP was a bit reduced and the K_D_ value of 0.24 μM was substantially higher than the K_D_ in the absence of c- di-AMP (compare Fig. 4A with 4B; Table 1). When *Bs*ComFB protein was saturated with 150 µM c-di-GMP, the binding of c-di-AMP was further reduced with a K_D_ of 121.1 µM (Table 1; compare Fig. 5A with 5B).

**Figure 4.**
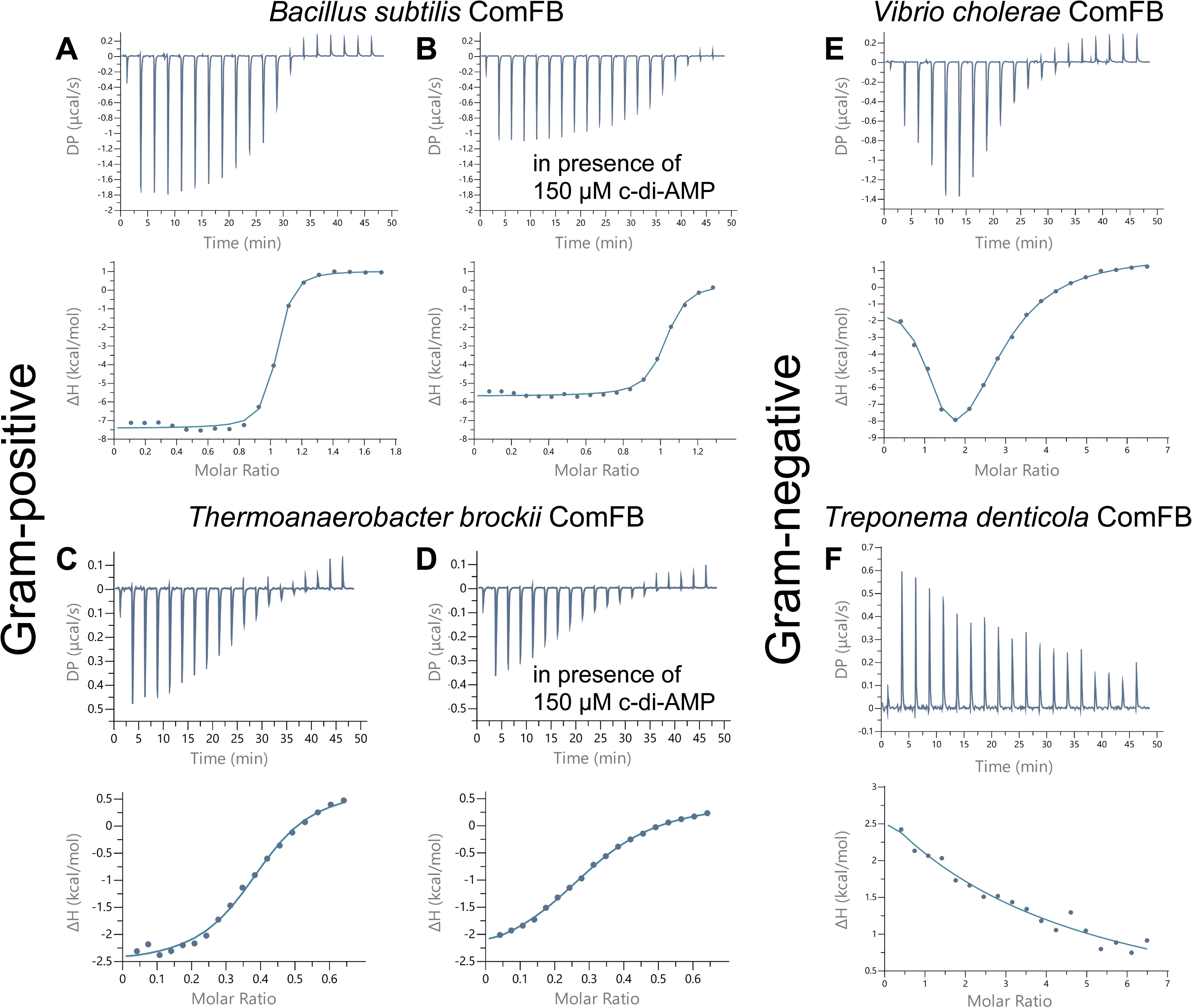
Isothermal titration calorimetry (ITC) analysis of c-di-GMP binding to phylogenetically different ComFB proteins. Upper panels show the raw ITC data in the form of heat produced during the titration of c-di-GMP on different ComFB proteins; lower panels show the binding isotherms and the best-fit curves according to the one binding site model. (A-D) ITC analysis of c-di-GMP binding to *B. subtilis* or *T. brockii* ComFB proteins in the absence (A,C) or presence of 150 µM c-di-AMP (B,D). (E,F) ITC analysis of c-di-GMP binding to *V. cholerae* (E) or *T. denticola* (F) ComFB proteins.

**Figure 5.**
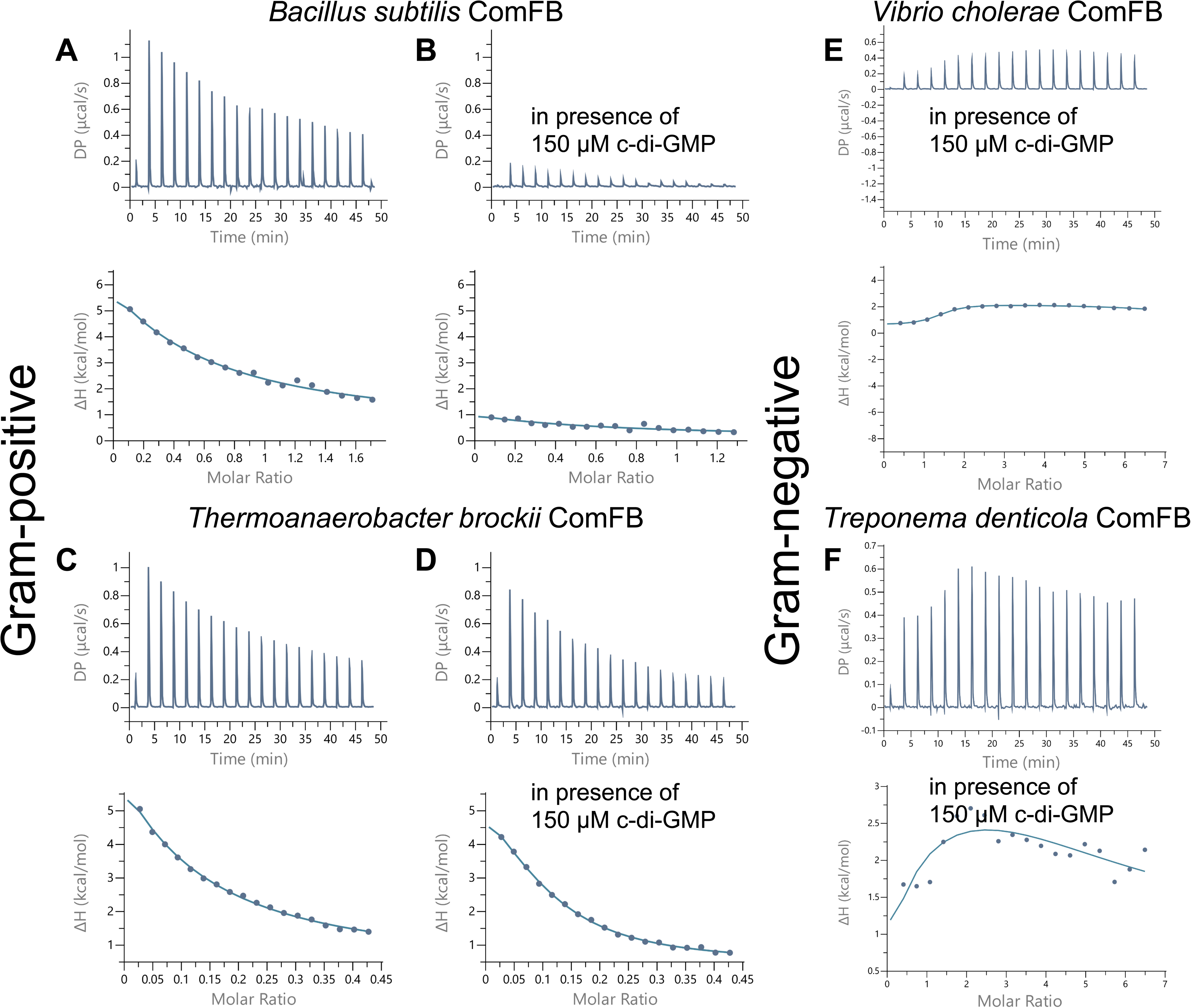
Isothermal titration calorimetry (ITC) analysis of c-di-AMP binding to phylogenetically different ComFB proteins. Upper panels show the raw ITC data in the form of heat produced during the titration of c-di-GMP on different ComFB proteins; lower panels show the binding isotherms and the best-fit curves according to the one binding site model. (A-D) ITC analysis of c-di-AMP binding to *B. subtilis* or *T. brockii* ComFB proteins in the absence (A,C) or presence of 150 µM c-di-GMP (B,D). (E,F) ITC analysis of c-di-AMP binding to *V. cholerae* (E) or *T. denticola* (F) ComFB proteins in the presence of 150 µM c-di-GMP.

**Table 1.**
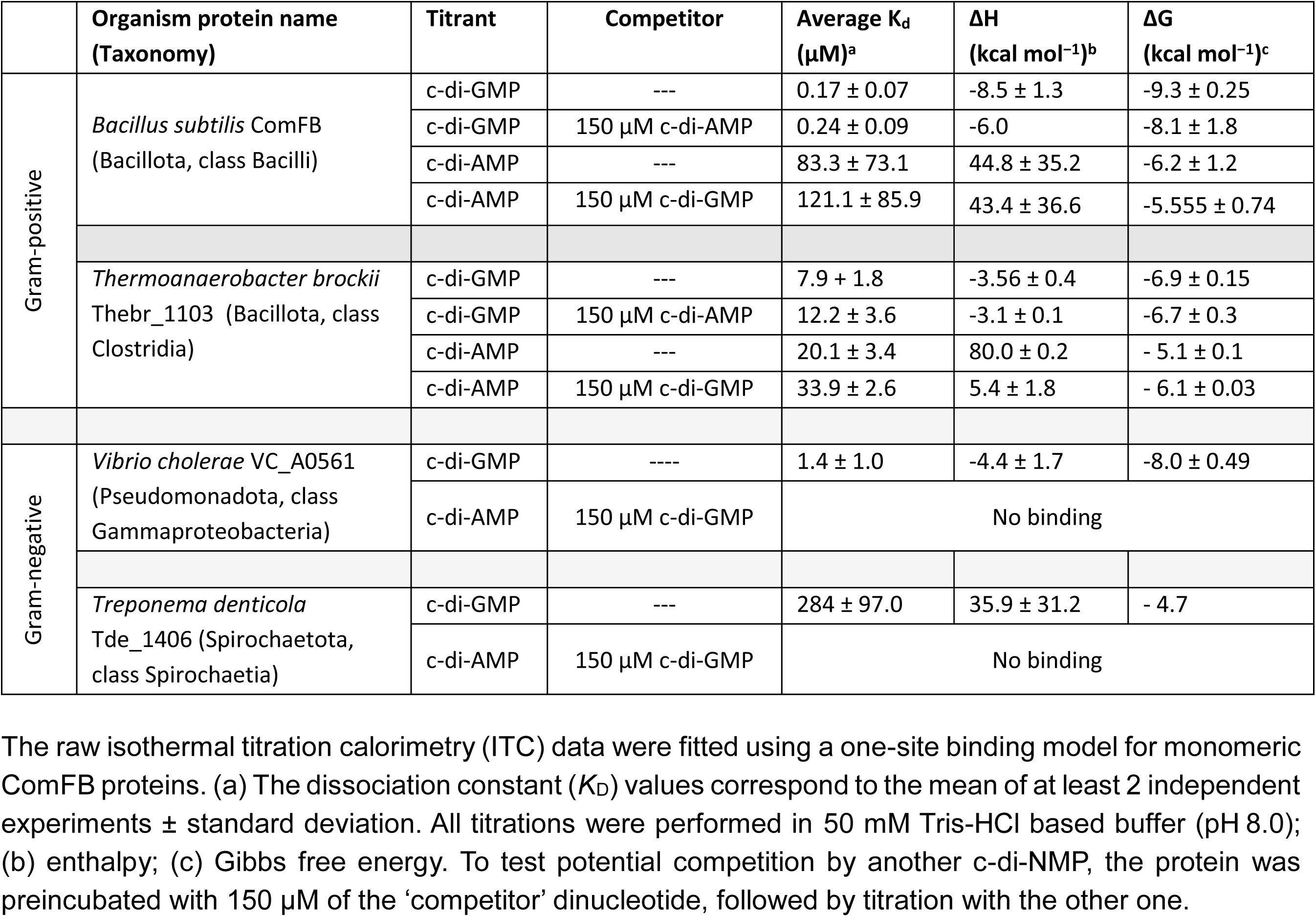
c-di-NMP binding by different ComFB proteins.

Again, the binding assays for ComFB from *T. brockii* (*Tb*ComFB) showed exothermic signals for c-di-GMP binding and endothermic profile for c-di-AMP binding (Fig. 4 and Fig. 5). However, compared to *Bs*ComFB, *Tb*ComFB showed weaker affinity towards c-di-GMP (Fig. 4C) with a K_D_ value of 7.9 μM and enhanced affinity towards c-di-AMP (Fig. 5C) with a K_D_ value of 20.1 μM (Table 1), implying that *Tb*ComFB could bind both c-di-GMP/c-di-AMP *in vivo*, similar to cyanobacterial *Sy*ComFB (Samir *et al*. 2023). The competition binding assays at saturating concentration of 150 µM of c-di-AMP (Fig. 4D) or c-di-GMP (Fig. 5D) also revealed that both molecules could compete with each other for binding to *Tb*ComFB. However, the binding affinity was substantially reduced compared to that in the absence of the competitor nucleotide (Table 1; compare isotherm of Fig. 4C with 4D and Fig. 5C with 5D).

Finally, we assessed the ability of the ComFB proteins from the gram-negative bacteria *V. cholerae* (*Vc*ComFB) and *T. denticola* (*Td*ComFB) to bind c-di-GMP or c-di-AMP. Remarkably, the *Td*ComFB possesses in addition to the N-terminal ComFB domain, a C-terminal immunoglobulin-like domain (Fig. S4). As expected, both *Vc*ComFB and *Td*ComFB proteins were able to bind to c-di-GMP (Table 1), however *Td*ComFB bound to c-di-GMP endothermically (Fig. 4F) with very low affinity (K_D_ 284 µM) compared to exothermic binding (Fig. 4E) and high affinity with K_D_ value of 1.4 µM for *Vc*ComFB. Surprisingly, both *Vc*ComFB and *Td*ComFB proteins were not able to bind c-di-AMP and c-di-AMP did not compete with c-di-GMP (Table 1 and Fig. 5E,F).

To further confirm that *Vc*ComFB binds only c-di-GMP, we used nano differential scanning fluorimetry (nanoDSF) compared to *Tb*ComFB, which binds both c-di-GMP and c-di-AMP with good affinity (Table 1). Both c-di-GMP and c-di-AMP thermally stabilized *Tb*ComFB, but only c-di-GMP stabilized *Vc*ComFB (Fig. S6), which further confirms that *Vc*ComFB binds only c-di-GMP, while *Tb*ComFB binds both c-di-NMPs. As all ComFB proteins showed robust binding of c-di-GMP and, in some cases, c-di-AMP (e.g. *Tb*ComFB and *Sy*ComFB), these experiments establish the ComFB superfamily as c-di-NMP receptor proteins, with preferential binding of c-di-GMP.

### Physiological importance of *comFB* in *Bacillus*

Next, to determine the biological significance of *comFB*, we investigated the *in vivo* function of *comFB* in the genetically accessible model bacterium, *B. subtilis*. As mentioned previously, *B. subtilis comFB* is localized in the vicinity of flagellar genes required for motility (Fig. 2), and it binds specifically to c-di-GMP (Fig. 4; Table 1), which is well known to inhibit motility in many bacteria, including *B. subtilis* (Wolfe & Visick, 2008, Römling *et al*., 2013, Chen *et al*., 2012, Subramanian *et al*., 2017). To examine whether ComFB plays a role in motility and if its function depends on c- di-GMP signaling, we expressed it from a constitutive promoter in the ectopic *amyE* locus of *B. subtilis* in the background of both wild-type and *ΔpdeH*, a phosphodiesterase mutant characterized by elevated levels of c-di-GMP (Chen *et al*., 2012, Gao *et al*., 2013). In these strains, growing in LB where competence is not expressed, the ComK-driven promoter in front of the *comF* operon is not active, and the main source of ComFB will be from the construct in *amyE.* Along with negative controls carrying empty vectors, all strains were inoculated in the centers of Petri dishes containing LB Medium with 0.3% agar and permitted to grow overnight. As shown in Fig. 6, ectopic expression of ComFB markedly inhibits swimming, and this inhibition is propagated in the absence of PdeH, the only c-di-GMP phosphodiesterase of *B. subtilis*. Deletion of *pdeH* significantly affects swimming by itself, presumably due to its effect via MotI (Subramanian *et al*., 2017). To rule out the sensitivity of the motility assay to growth differences, we additionally compared the growth of the relevant strains and detected no significant differences (Fig. S7). These data provide strong evidence that ComFB interferes with motility in the presence of elevated c-di-GMP, confirming the biological relevance of its high affinity for c-di-GMP (Table 1).

**Figure 6.**
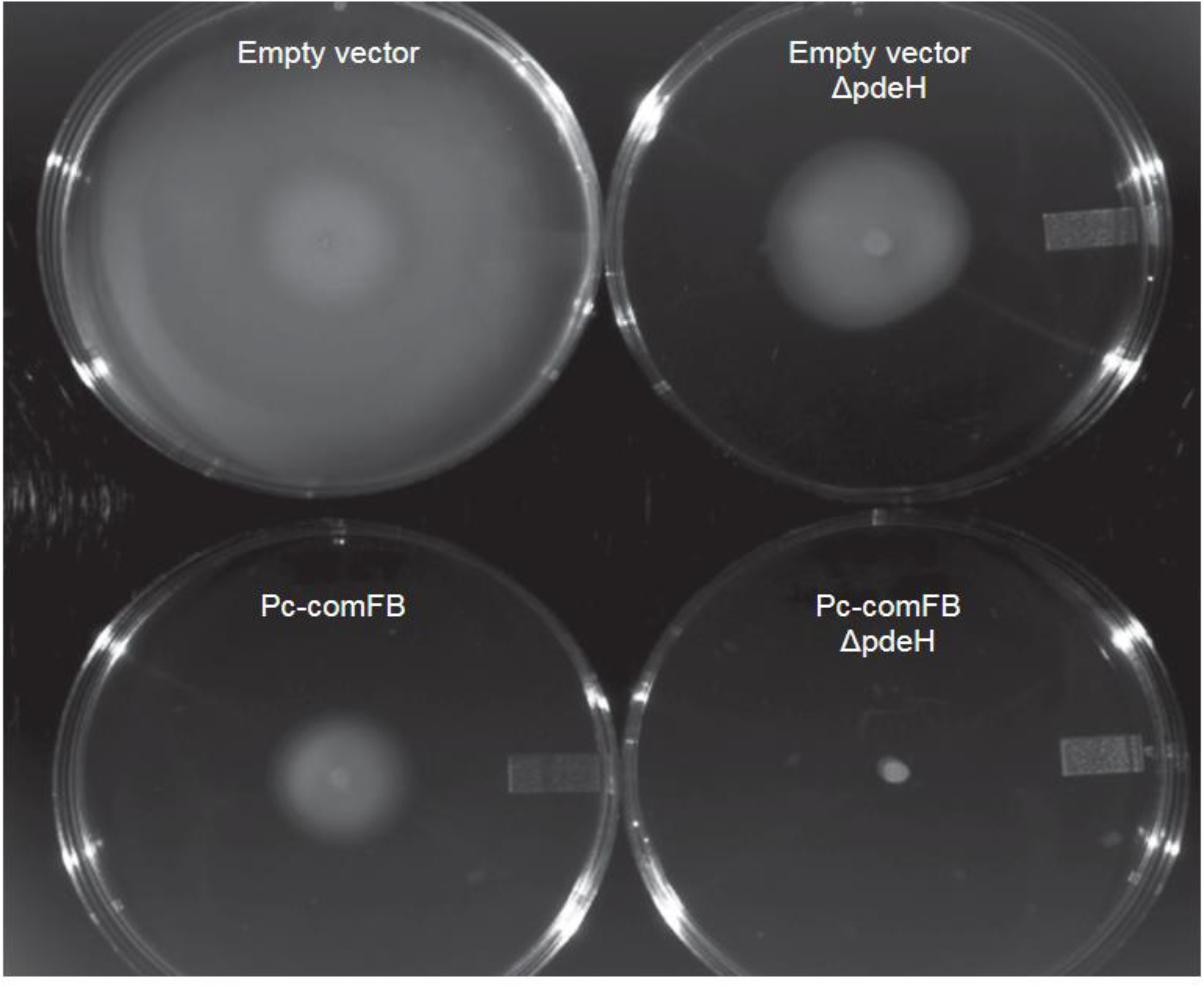
ComFB inhibits swimming. The swimming assay was conducted as described in Methods by inoculating cells into 0.3% agar LB plates. Plasmid vectors carrying *comFB* under the control of a constitutive promoter or the same vector without comFB (empty vector) were inserted separately at *amyE* in wild type and *ΔpdeH* backgrounds. The image was acquired after 20 hours of growth at 30 C in a humidified chamber, followed by a further 5 hours at 37 C.

## DISCUSSION

Proteins of the ComFB family (Pfam domain <u>PF10719</u>) are widespread in bacteria, being encoded, besides *Bacillota* and *Cyanobacteriota*, in the genomes of representatives of at least five other phyla (Fig. 1B and Fig. S1). Further, metagenomic sequencing identified *comFB* genes in members of a dozen more phyla (Table S1) and several candidate phyla. These proteins are usually encoded in a single copy per genome, such as CdgR from *Nostoc* sp. PCC 7120, but are occasionally found in multiple copies, from two genes in *Alkaliphilus metalliredigens* (Amet_2487 and Amet_3088) to six paralogous copies in the genomes of *Synechococcus* sp. PCC 7502 and *Pseudanabaena* PCC 7367 and seven in *Allocoleopsis franciscana* PCC 7113 (Table S1).

With respect to cyanobacterial c-di-GMP signaling, the discovery of CdgR (Zeng *et al*., 2023) closed an important gap in knowledge: the paucity of known c-di-GMP receptors in cyanobacteria. Indeed, as seen in the c-di-GMP census, *Nostoc* sp. PCC 7120 and *Synechocystis* sp. PCC 6803 encode multiple diguanylate cyclases (Enomoto *et al*., 2023), but no PilZ-domain proteins and very few MshEN-containing proteins (1 and 3, respectively). The c-di-GMP binding by these MshEN domains has not been tested so far, although c-di-GMP has been shown to control cyanobacterial cell motility and phototaxis (Savakis *et al*., 2012, Angerer *et al*., 2017, Enomoto *et al*., 2023, Wallner *et al*., 2020). Other cyanobacteria encode PilZ domains as part of their bacterial cellulose synthase (*bcsA*) genes and MshEN domains at the N-termini of their PilB/GspE ATPases, which control twitching motility and type II secretion. No other cyanobacterial c-di-GMP targets had been characterized until last year. Characterization of CdgR, which is widespread in cyanobacteria, and demonstration of its involvement in controlling cell size (Zeng *et al*., 2023) provided a much-needed rationale for the presence of multiple c-di-GMP turnover enzymes in various cyanobacterial genomes.

While Zeng and colleagues assumed that CdgR was specific for cyanobacteria (Zeng *et al*., 2023), the structural similarity between CdgR and ComFB (Fig. 1A), coupled with the conservation of several c-di-GMP-binding residues within the entire ComFB/CdgR family (Fig. 1B), suggested that other members of this family might also bind c-di-GMP, such that this family would represent a widespread c-di-GMP receptor protein. This suggestion was verified here by demonstrating high-affinity c-di-GMP binding by *Bs*ComFB (Table 1), the founding member of the family, whose proposed role in competence had been used for annotating the entire family (<u>PF10719</u> in Pfam database and <u>IPR019657</u> in InterPro) (Mistry *et al*., 2021, Paysan-Lafosse *et al*., 2023) as “late competence development protein ComFB”. *Bs*ComFB was shown to bind c-di-GMP with high affinity, Kd = 0.17 µM, which is remarkably close to the value previously reported for the CdgR from *Nostoc* sp. PCC 7120 (Zeng *et al*., 2023).

Further, three other ComFB family members from diverse bacterial lineages were also found to bind c-di-GMP (Table 1). The shared ability to bind c-di-GMP by ComFB-like proteins from *B. subtilis*, two cyanobacteria, and these three organisms strongly suggests that the ComFB family represents the missing widespread family of c-di-GMP receptors, the third one after PilZ and MshEN domains. Accordingly, the entire family can now be renamed “ComFB-like c-di-GMP receptors.”

The new name also makes sense because of the uncertain role of *Bs*ComFB in competence. Genome analysis has revealed the absence of the *comFB* gene in several *Bacillus* species that appeared to be transformable (Kovacs *et al*., 2009, Kovacs *et al*., 2013). Further, while *comFA- comFC* operons are fairly widespread, they are commonly found in the genomes that do not encode *comFB* (Table S2); the *comFA-comFB-comFC* operons, like the one in *B. subtilis* are only found in a few closely related *Bacillus* species. Finally, a *B. subtilis* strain expressing *comFA* and *comFC* in the absence of *comFB* was reported to be normally transformable (Sysoeva *et al*., 2015). These data suggested that ComFB may not be required for competence in *B. subtilis*.

Nevertheless, the ability of ComFB proteins to bind c-di-GMP, the master regulator of motility-related functions, and the genomic association of *comFB* with flagella (as seen in *B. subtilis* and *T. denticola*) or pili (as seen in cyanobacteria, *Caldicellulosiruptor* and *Thermoanaerobacterium* species) genes still suggests a possible involvement of ComFB proteins in regulation of cell motility. However, the role of ComFB might be different in other organisms. In this context, we were able to show that ComFB is participating in a c-di-GMP-mediated inhibition of motility in *B. subtilis* using swimming assay (Fig. 6). However, it is important to note that the swimming assay does not distinguish between an effect of ComFB on the production or function of flagella, not does it exclude an effect on chemotaxis. It will require further investigation to distinguish between these alternatives. Nevertheless, *Bs*ComFB joins MotI (Subramanian *et al*., 2017) as a c-di-GMP binding protein associated with a motility-related phenotype in *B. subtilis*. Moreover, our recent work showed that *Sy*ComFB (encoded by *slr1970*), the CdgR ortholog from the model cyanobacterium *Synechocystis* sp. PCC 6803, is required for pilus biogenesis and natural competence, albeit through its interaction with c-di-AMP, another bacterial second messenger (Samir *et al*., 2023). These data, along with the recently demonstrated roles of CdgR in the regulation of cell size (Zeng *et al*., 2023), cyanobacterial motility, and DNA uptake, highlight the unique properties of the ComFB family proteins as dedicated receptors for both c-di-NMP second messengers, c-di-GMP and c-di-AMP.

The demonstration of c-di-GMP-binding by ComFB family proteins leaves many questions to be addressed in future studies. One of them is the ability of (some) ComFB proteins to bind a metal atom, such as Zn^2+^, and, if so, the potential regulatory role(s) of this metal atom. The Zn^2+^-binding Cys residues of *Bs*ComFB are conserved in other members of the phylum *Bacillota*, as well as in Deferribacterota, Thermotogota, and some spirochetes (Fig. S1), suggesting that these proteins may also contain Zn^2+^ atoms. The presence of metal atoms in the ComFB proteins from Betaproteobacteria and Gammaproteobacteria, which contain a single conserved Cys residue (Fig. S1) is uncertain. Some alteromonads, however, have an additional Cys residue in the Cys-Cys motif (Fig. S1). Accordingly, a metalloproteome study of the marine bacterium *Alteromonas* sp. BB2-AT2 found that its ComFB contained Zn^2+^ ions (Mazzotta *et al*., 2021). Anyway, given the structural similarity and similar c-di-GMP binding properties of Zn^2+^-containing *Bs*ComFB and Zn^2+^-less CdgR (Fig. 1A), the structural role of Zn^2+^ atoms is likely to be dispensable in the ComFB family proteins but might play a role in the regulation of their binding affinities.

In conclusion, this work, along with the recent finding on cyanobacterial *comFB* (Zeng *et al*., 2023, Samir *et al*., 2023), shows that diverse members of the ComFB family serve as dedicated receptors for c-di-GMP and/or c-di-AMP and play important regulatory roles in diverse lineages of bacteria. The ability of at least some ComFB proteins to bind two distinct second messengers and the diversity of *comFB*-containing operons (Fig. 2) and ComFB domain architectures (e.g., in combination with PATAN, REC, CheR, PspA/IM30, immunoglobulin-like, or other domains; Fig. 3 and Fig. S3), suggest that the spectrum of these regulatory roles could be quite wide. This establishes the ComFB superfamily as new c-di-NMP receptor proteins widespread in almost all bacterial domains.

## MATERIALS AND METHODS

### Sequence and structure analysis

Structural alignment of the ComFB protein from *B. subtilis* (GenBank: <u>CAB15563</u>; PDB: <u>4WAI</u>) and *Sy*ComFB (Slr1970) from *Synechocystis* sp. PCC 6803 (GenBank: <u>BAA18199</u>; PDB: <u>8HJA</u>) was performed with Dali (Holm, 2022). Secondary structure elements were taken from the Dali alignment and adjusted using DSSP outputs of the PDB data. The alignment was visualized with PyMol (Schrödinger, LLC). Closely and distantly related members of the ComFB/CdgR superfamily were retrieved through iterative searches of the NCBI protein database using PSI-BLAST (Altschul *et al*., 1997) and of UniProt using HMMer (Potter *et al*., 2018), respectively. In addition, InterPro (Paysan-Lafosse *et al*., 2023) was used to retrieve ComFB family members assigned to the Pfam domain <u>PF10719</u> (InterPro entry <u>IPR019657</u>). Additional proteins, such as *Anabaena cylindrica* Anacy_5104 (UniProt: K9ZNS1) or *Trichormus variabilis* Ava_3600 (UniProt: Q3M730), that were annotated as ComFB in UniProt but not included in these InterPro and Pfam families, were verified with HHpred (Zimmermann *et al*., 2018) and then used as queries in further PSI-BLAST and HMMer searches. The sequences obtained (see Fig. S1 and Table S1) were added to the ComFB superfamily.

The presence or absence of ComFB family members in certain bacterial lineages, as well as ComFB domain counts in individual bacterial genomes, were evaluated using taxonomy-restricted BLAST searches and taxonomy-based sorting of the BLAST outputs. Sequence alignments of distant family members against *B. subtilis* ComFB and *Sy*ComFB (Slr1970) structures were verified using HHpred (Zimmermann *et al*., 2018) tool of the MPI Bioinformatics Toolkit (Gabler *et al*., 2020). Domain architectures of the ComFB proteins were analyzed using the CDD (Wang *et al*., 2023) and Pfam (Mistry *et al*., 2021) databases; domain assignments were checked with HHpred (Zimmermann *et al*., 2018).

### Cluster map analysis

To gather protein sequences for cluster analysis, we queried the UniProt database for homologs of the ComFB protein sequences from *B. subtilis* (P39146), *Synechocystis* sp. PCC 6803 (P74113), *Vibrio cholerae* (Q9KM28), and *Fischerella muscicola* (A0A2N6JYB2, A0A2N6JYC8) using BLAST with an E-value threshold of 1e-04 and a “max_target_seqs” parameter of 20,000. We aggregated the full-length sequences of the resulting matches, removing any flagged as “Fragment,” and then filtered them using MMseqs2 to retain sequences with a maximum pairwise identity of 70% and a length coverage of at least 70%. The resulting filtered set, comprising a total of 1,626 sequences, was subjected to clustering analysis using CLANS (CLuster ANalysis of Sequences) tool (Frickey & Lupas, 2004) based on all-against-all pairwise P-values calculated with BLAST. The clustering was performed until equilibrium was reached in a 2D space, applying a P-value cutoff of 1e-06 and using the default settings in CLANS. The clustering was visualized using CLANS.

### Protein production and purification

The plasmids and primers used in this study are listed in Table S3. The plasmids expressing *comFB* of *Bacillus subtilis* (BSU_35460, GenBank: <u>CAB15563</u>) and *Vibrio cholerae* (VC_A0561, GenBank: AAF96463.1) were constructed using polymerase chain reaction (PCR) using a genomic DNA *B. subtilis* and *V. cholerae* as a template, while *comFB* of *Thermoanaerobacter brockii* (Thebr_1103, GenBank: ADV79679.1) and *Treponema denticola* (Tde_1406: GenBank: AAS11923.1) were constructed using a designed gBlocks DNA fragments (IDT, USA) as described previously (Selim *et al*., 2018). The *comFB* genes were cloned into a pET-28a vector with Gibson assembly, incorporating a C-terminal 8xHis-tag. Positive clones were selected on agar plates supplemented with 50 μg/ml kanamycin. The recombinant proteins were expressed and purified as described previously in *E. coli* strain LEMO21 (DE) (Selim *et al*., 2021b, Selim *et al*., 2019). Briefly, the 8xHis-tagged constructs were expressed by overnight cultivation of *E. coli* cells at 20°C in the presence of 0.5 mM IPTG and purified by immobilized metal affinity chromatography using Ni^2+^-Sepharose resin (Cytiva^TM^), followed by size exclusion chromatography on a Superdex 200 Increase 10/300 GL column (GE HealthCare, Munich, Germany), as described previously (Selim *et al*., 2020). Protein purity was assessed by Coomassie-stained SDS-PAGE, and protein concentrations were determined using Bradford assay.

### Size exclusion chromatography and multiangle light scattering analysis (SEC-MALS)

Analytical size exclusion chromatography was carried out at room temperature, as described previously, using the ÄKTA purifier (GE Healthcare) on Superdex^TM^ ^200^ prep-grade column (GE Healthcare) coupled to a multiangle light scattering (MALS) detector (Selim *et al*., 2019, 2020; (Walter *et al*., 2019)). The protein samples diluted 1:4, centrifuged for 5 min at 15000 g, and 250 µl of the supernatant were injected into the column with a flow rate of 0.4 ml/min, after equilibrating the column with the running buffer (50 mM Tris/HCl, pH 8.0, 300 mM NaCl). ASTRA software (Wyatt) was used for data analysis and molecular mass calculations using the MALS data. The elution volume was plotted against the UV signal and molecular mass (Selim *et al*., 2019).

### Isothermal titration calorimetry (ITC) analysis and nucleotides binding assays

Binding of c-di-GMP or c-di-AMP by recombinantly produced ComFB proteins was analyzed by isothermal titration calorimetry (ITC), as described previously (Haffner *et al*., 2023a, Mantovani *et al*., 2024). Protein samples for ITC were dialyzed against the assay buffer (50 mM Tris/HCl, pH 8.0, 300 mM NaCl, 0.55 mM EDTA), then diluted with the same buffer to a working concentration of 30-300 µM, before loading into the ITC cell. C-di-GMP/c-di-AMP was dissolved in the same buffer to the concentration of 1.0 mM and added gradually in 2 µl injections using a microsyringe. ITC measurements were conducted on a MicroCal PEAQ-ITC instrument (Malvern Panalytical, Westborough, MA, USA), at 25 °C, with a reference power of 10 μcal/s. Control experiments to determine the dilution heat of c-di-GMP or c-di-AMP were performed by titrating c-di-GMP or c- di-AMP into a cell filled with assay buffer. Dissociation constants K_D_ and ΔH values were calculated using the single binding site model with the MicroCal PEAQ-ITC Analysis Software (Malvern Panalytical) after subtracting the dilution heat of the control experiments. In the competition assay of c-di-GMP and c-di-AMP binding to ComFB, the protein was incubated with 150 µM of one of the nucleotides and titrated against 1 mM of the other competing ligand. For reproducibility, different batches of purified ComFB proteins were used for different ITC experiments. The details of ITC measurements are presented in Table 1 and Figures 4 and 5.

### Growth of bacteria and strain construction

Constructs were introduced into *B. subtilis* using the transformation of competent cells. Antibiotic selections were on LB-agar plates using spectinomycin (100 μg/ml) or kanamycin (5 μg/ml). All constructs were confirmed by sequencing. The *ΔpdeH::kan* construct and plasmid pKB149, which carries a constitutive promoter and confers spectinomycin resistance, were kind gifts from Dan Kearns (Indiana University) (Subramanian *et al*., 2017). For overexpression, the coding sequence of *comFB* was isolated by PCR using the P_c_-forward and Pc-Reverse primers (Table S3). The resulting DNA fragment was inserted into pKB149 after cutting with *Hind*III and *BamH*I, placing comFB under the control of a constitutive promoter (P_c_). The resulting plasmid was linearized using *Sca*I to favor replacement recombination and transformed into *B. subtilis* IS75 for insertion into the ectopic *amyE* locus. Correct insertion was confirmed by testing on starch-plates and the overexpression of ComFB was verified by Western blotting, using an antiserum raised against purified ComFB. The empty vector (pKB149) was similarly inserted in *amyE* to produce a control strain.

### Swim test

The method used was as described in (Hall *et al*., 2018). Single colonies were picked with toothpicks and inoculated into LB fortified with 0.35% agar. The plates were grown overnight at 30 C in a humidified incubator, then transferred to 37 C for further growth, and photographed when the control strain (BD9422) had nearly reached the outer edge of its Petri dish.

## ACKNOWLEDGEMENTS

The KAS laboratory is funded by grants from the German Research Foundation (DFG) as part of the priority research program (SPP2389; SE 3449/1-1) and by the collaborative research center SFB1381 (project number: 403222702). KAS also gratefully acknowledges the infrastructural support and funding by the Cluster of Excellence “Controlling Microbes to Fight Infections (CMFI)” (EXC2124–390838134) and by the Excellence Strategy of the German Federal and Baden-Württemberg State Governments (Projektförderung: PRO-SELIM-2022-14). MYG was supported by the Intramural Research Program of the National Library of Medicine at the U.S. National Institutes of Health. We thank Shan-Ho-Chou (Huazhong Agricultural University) for the initial version of (Fig. 1A), Karl Forchhammer (Tübingen University), Jörg Stülke (Göttingen University) and Annegret Wilde (Freiburg University) for continued support and constructive discussions. We are also grateful to Marcus Hartmann (MPI for Biology Tübingen), and to Heinz Grenzendorf and Filipp Oesterhelt (Tübingen University) for their excellent support and/or assistance. NIH Grant R01GM123487 supported the work in Dubnau lab. DD acknowledges valuable discussions with Dan Kearns, his expert advice, his generous gift of strains, and additional discussions with Louisa Celma and Jeanie Dubnau.

## Supplementary Materials

**Table S1**. A list of representative ComFB family members in various bacterial phyla.

**Table S2**. Phylogenetic distribution of ComFB, ComFA and ComFC proteins, according to the COG database.

**Table S3**. Plasmids and primers used in this study.

**Figure S1.**
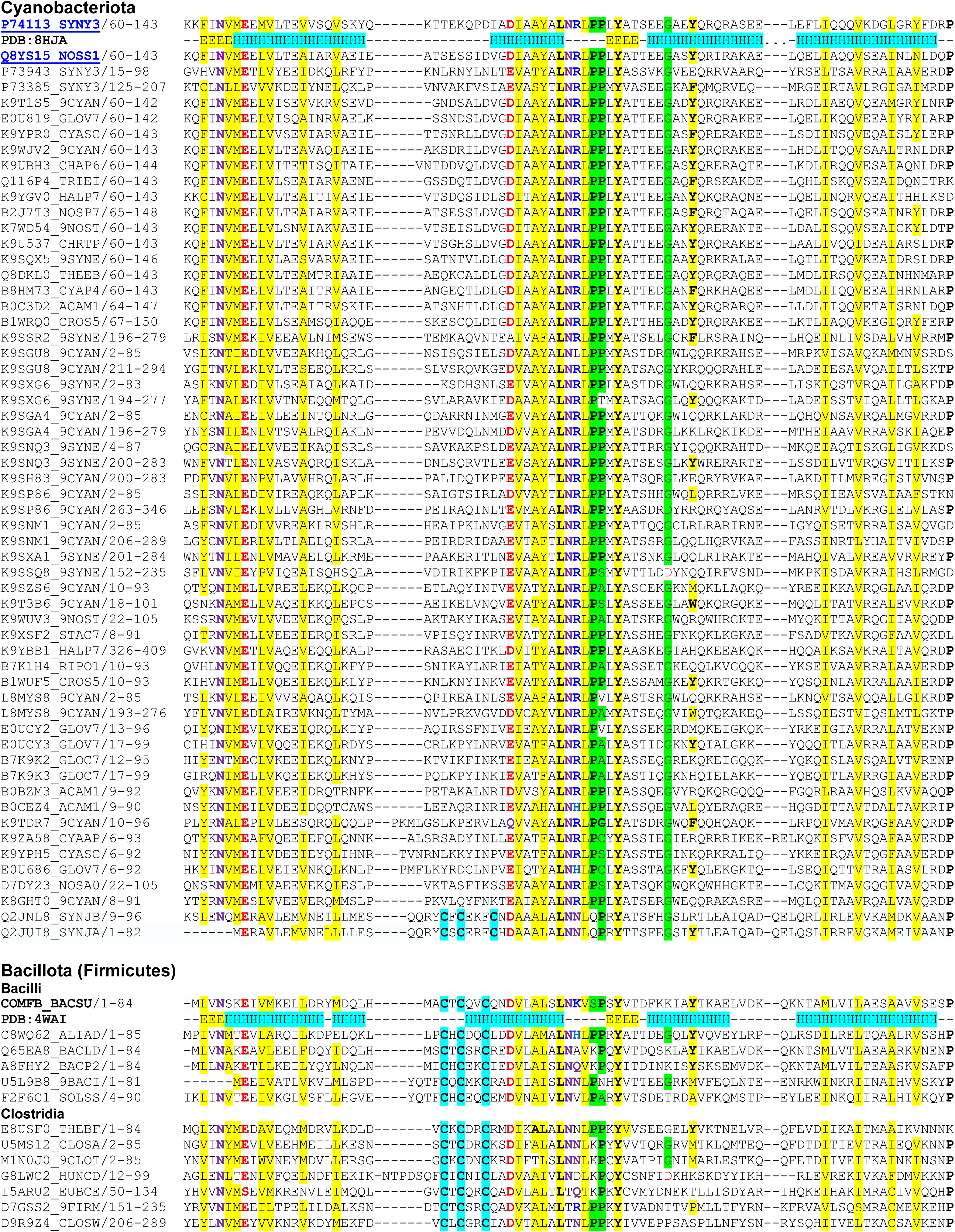
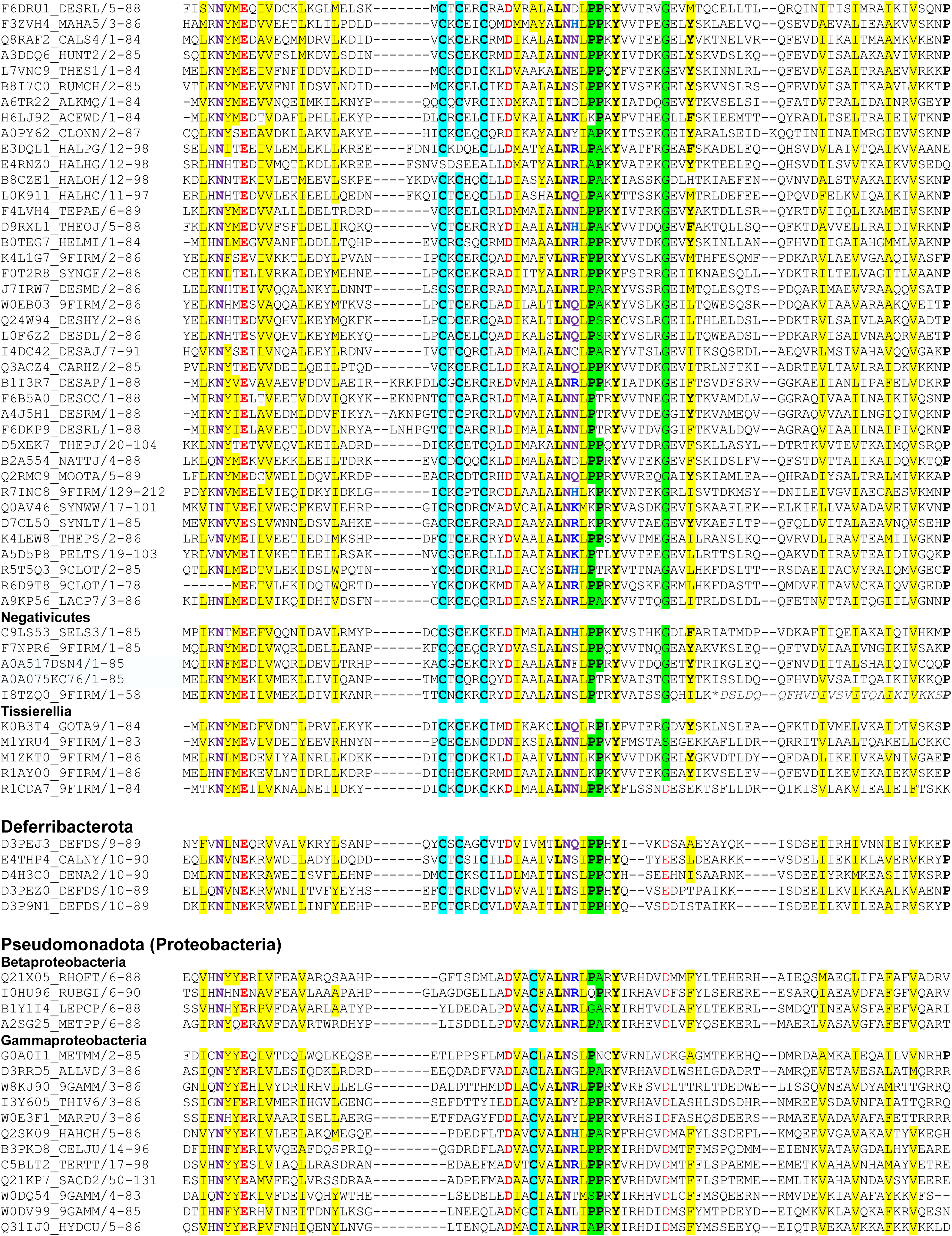
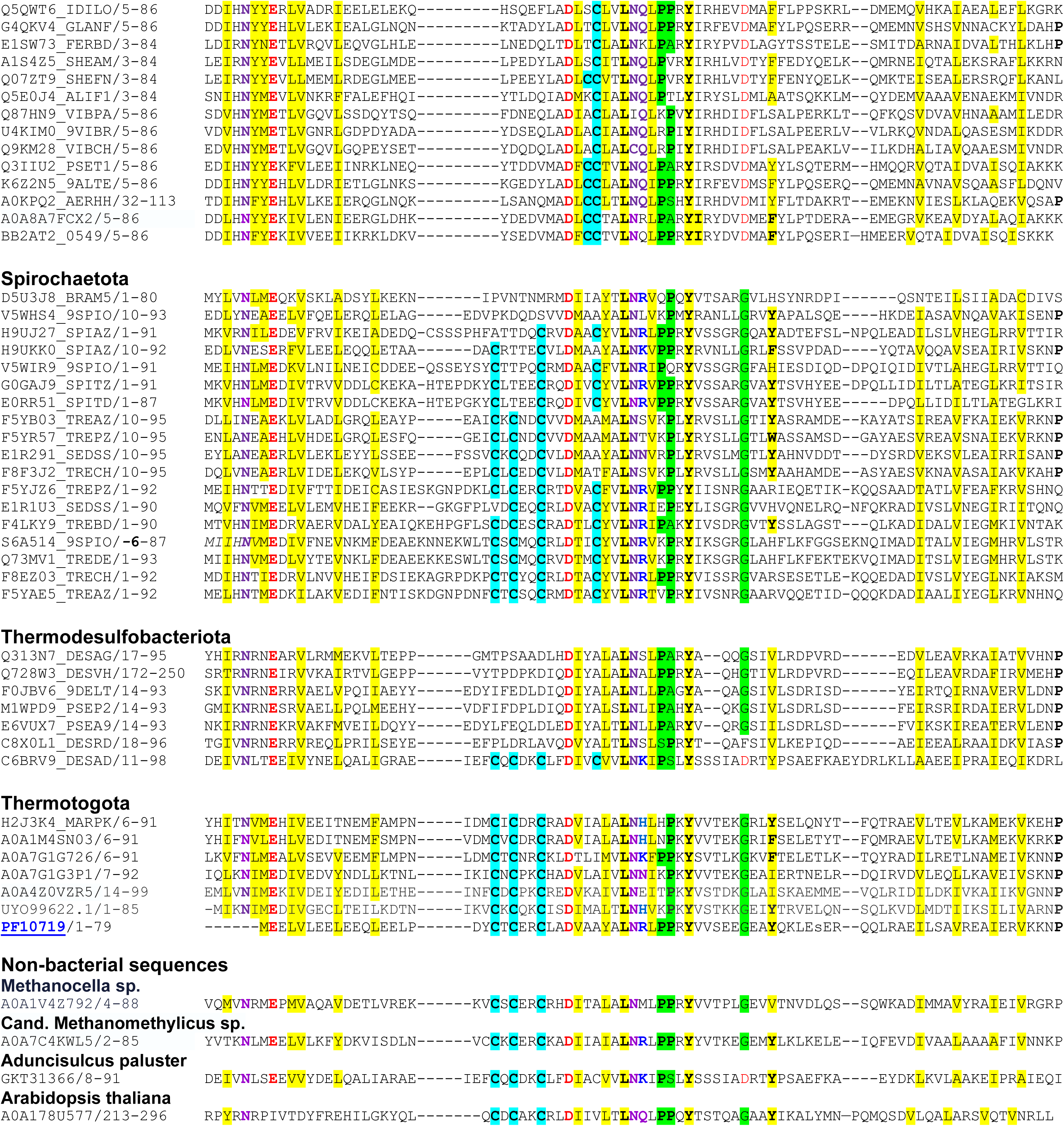
Sequence alignment of representative ComFB/CdgR family members

**Figure S2.**
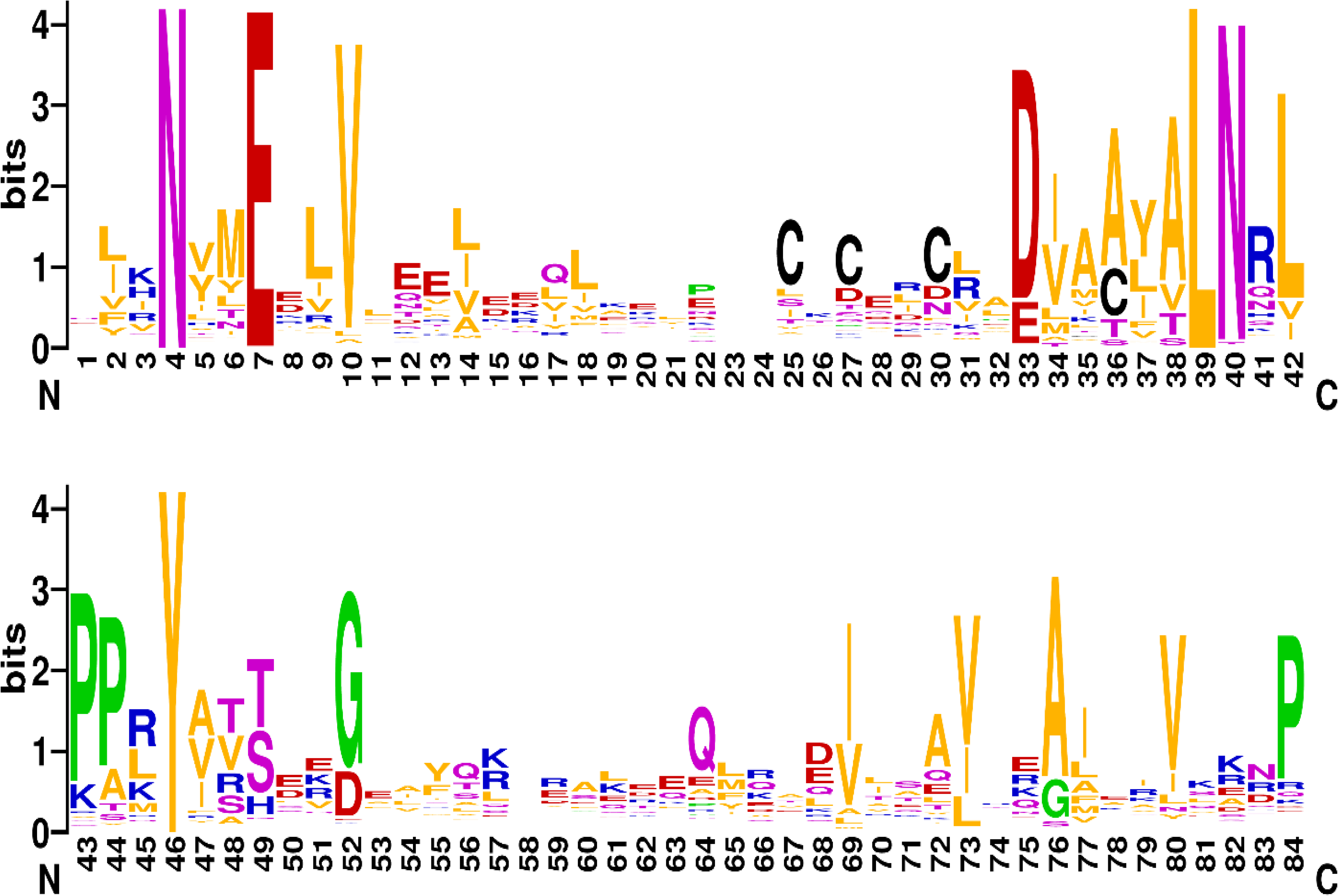
Sequence logo of ComFB/CdgR family, based on an alignment of 180 distinct family members.

**Figure S3.**
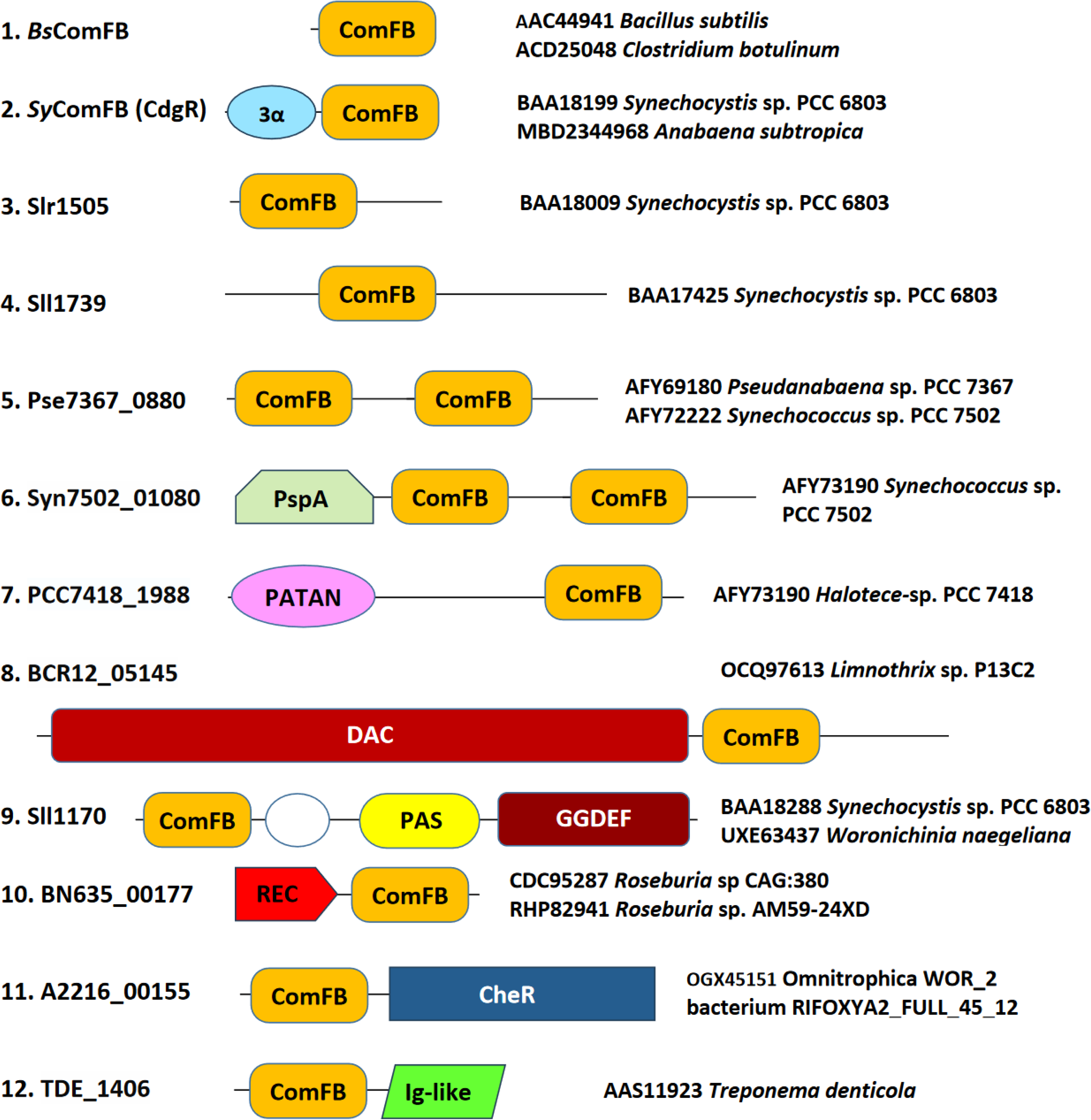
Domain architectures of various ComFB proteins.

**Figure S4.**
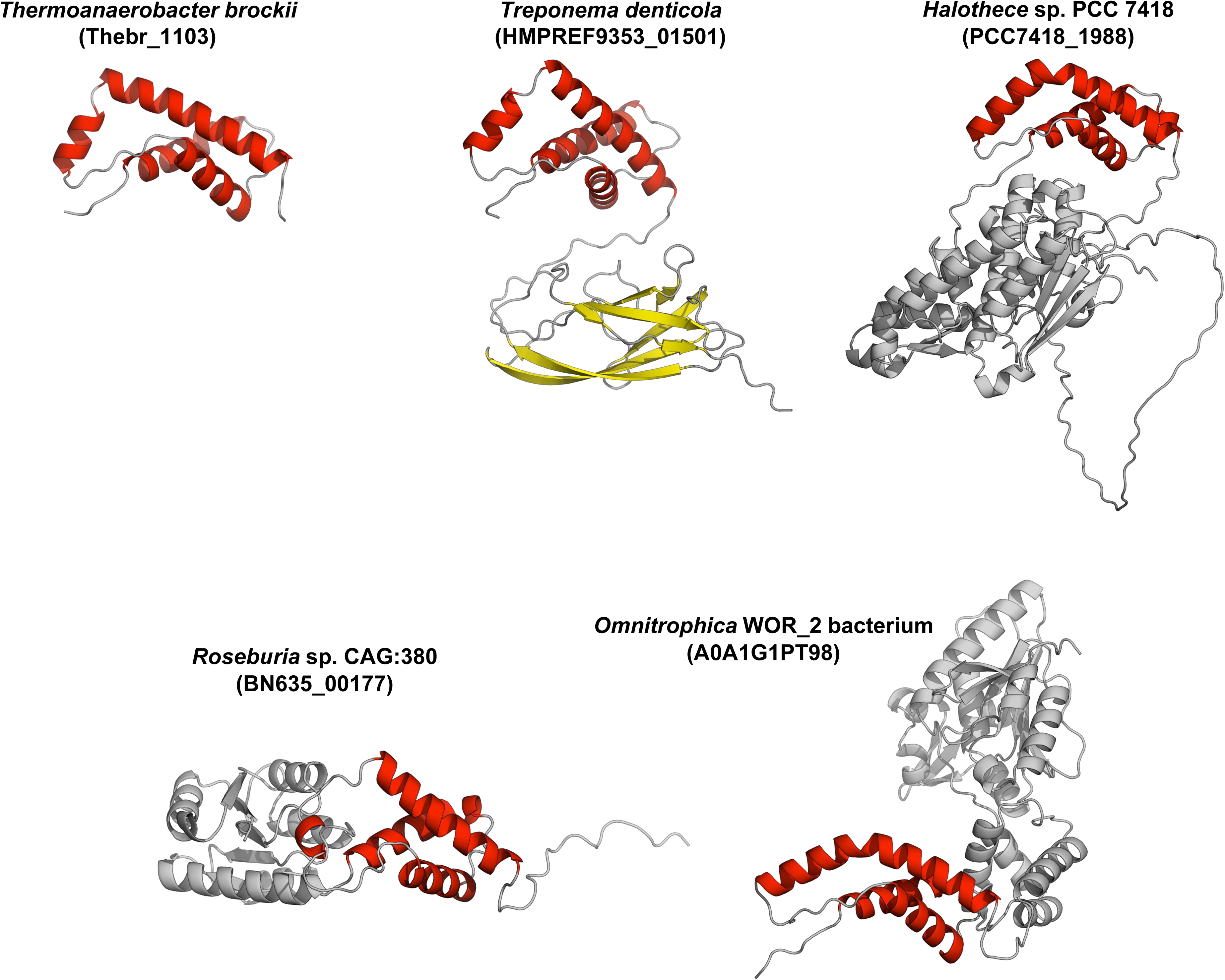
Additional representative structures of ComFB domain-containing proteins from various species. The structures are AlphaFold2 predictions obtained from the UniProt/AlphaFold DB, with the exception of *Treponema denticola*, which was predicted independently using AlphaFold2. The species represented include *Thermoanaerobacter brockii* (UniProt E8USF0), *Halothece* sp. PCC 7418 (K9YBB1), *Roseburia* sp. CAG:380 (R6WME1), and *Omnitrophica* WOR_2 bacterium (A0A1G1PT98). Four of these proteins feature additional domains: *T. denticola* contains an Ig-like domain, *Halothece* sp. PCC 7418 a PATAN domain, *Roseburia* sp. CAG:380 a REC domain, and *Omnitrophica* WOR_2 bacterium a CheR domain.

**Figure S5.**
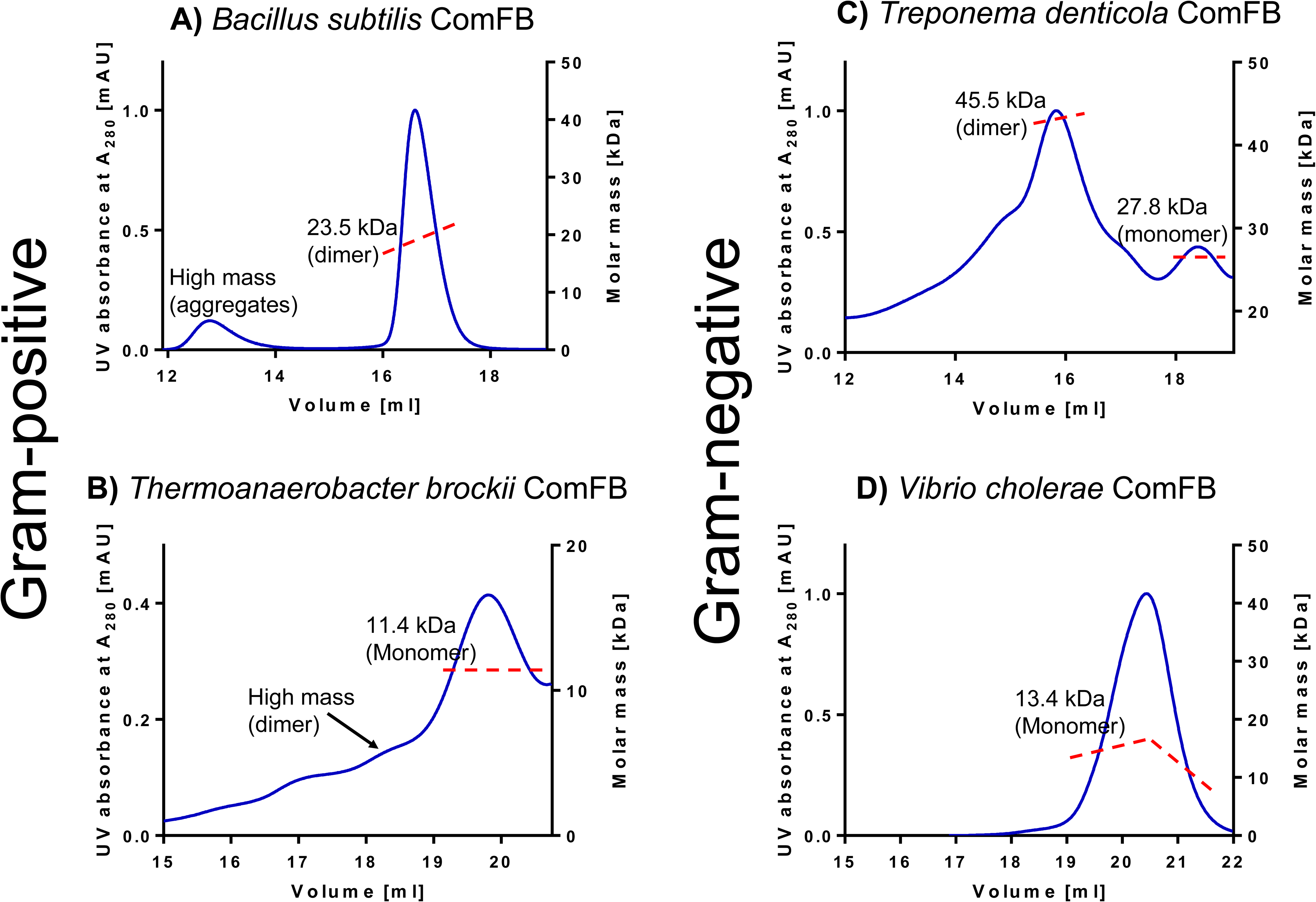
The oligomeric state of different ComFB proteins was determined by SEC-MALS.

**Figure S6.**
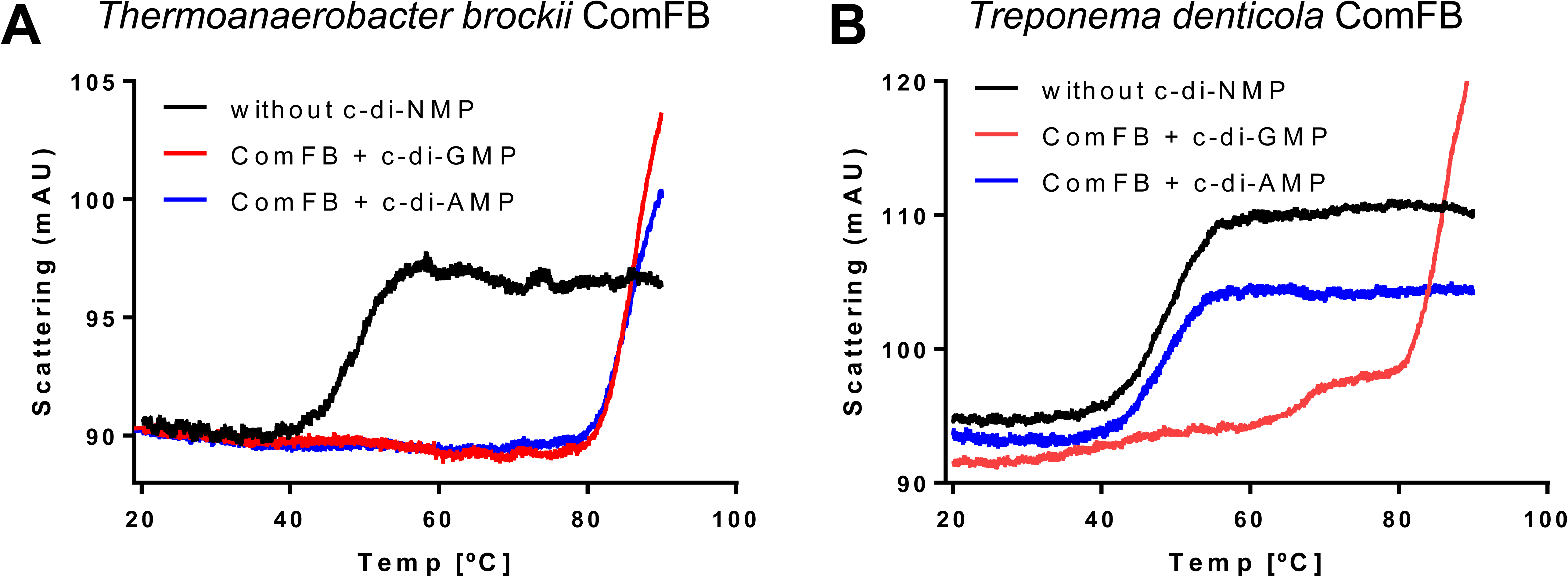
NanoDSF of c-di-NMP binding to ComFB proteins from *T. brockii* and *T. denticola.* Light scattering thermograms were obtained by measuring the attenuation of the back-reflected light intensity passing through the protein sample with or without 0.5 mM c-di-AMP or c-di-GMP as a function of temperature.

**Figure S7.**
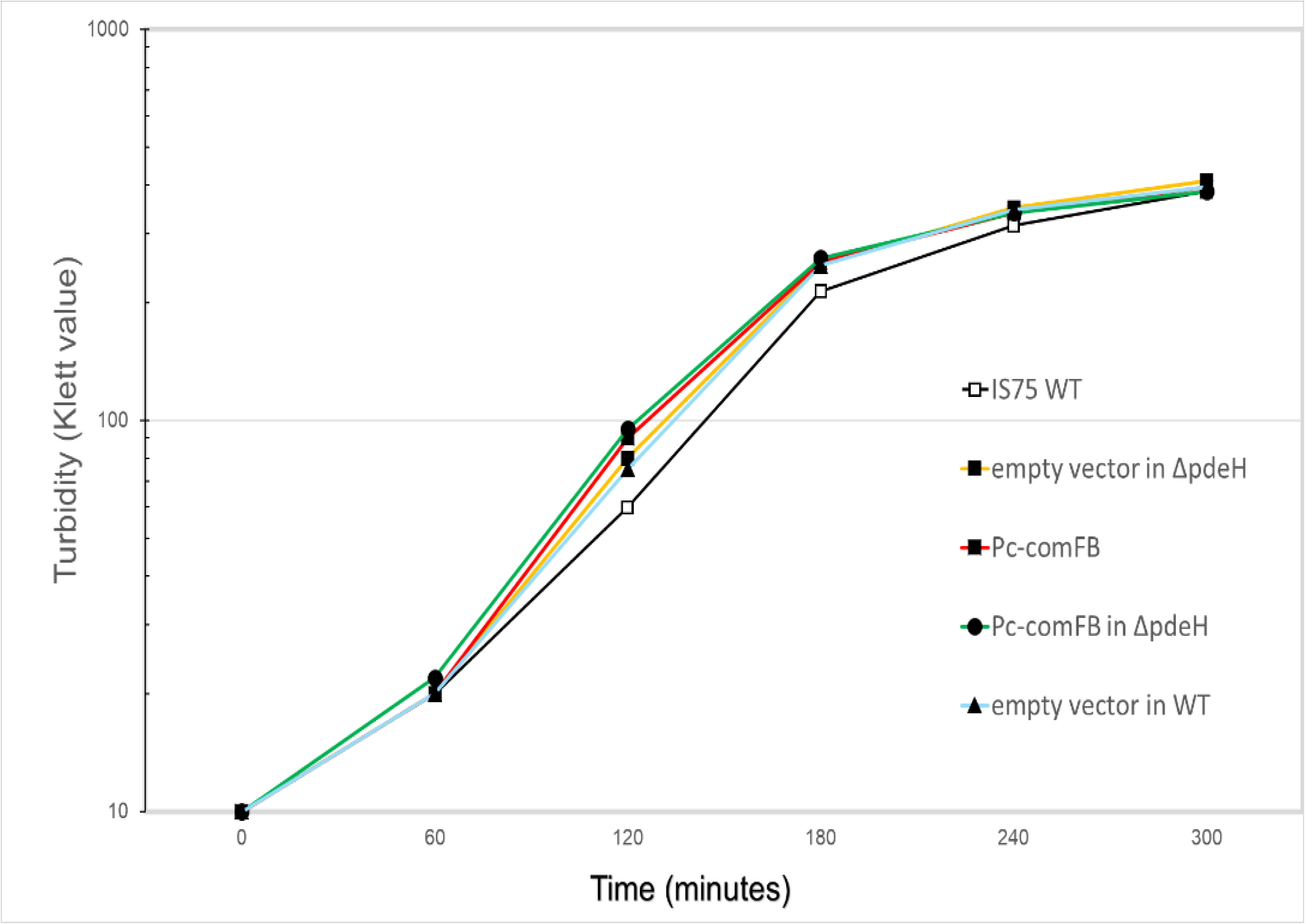
BD9422 (*amyE::*pDR511), BD9385 (*amyE::*pDR511 *pdeH::kan*), BD9389 (Pc-*comFB*), BD9398 (Pc-*comFB pdeH::kan*) and the wild-type strain IS75, were grown in liquid LB medium with vigorous shaking at 37 C. Growth was followed as turbidity with a Klett colorimeter.

