## Supplemental Table S3 for "ComFB, a new widespread family of c-di-NMP receptor proteins"

**Table S3**. Plasmids and primers used in this study

| **Recombinant proteins and plasmids** | **Primers/Sequence (5′→3′)** | **Note/ Refs.** |
|---|---|---|
| $His_8$-tagged *Bs*ComFB (pET28a_*Bs*ComFB-$His_8$ plasmid) | Bs_ComFB-His_Fw:<br>CTAGAAATAATTTTGTTTAACTTTAAGAAGGAGATATACCATGCTTGTCAATTCAAAAGAAATCG | This study |
| | Bs_ComFB-His_Rev:<br>CAGCCGGATCTCAGTGGTGGTGGTGGTGGTGGTGGTGCGAGTTTATGAATGCCTCCTCCTGC | |
| $His_8$-tagged *Vc*ComFB (pET28a_*Vc*ComFB-$His_8$ plasmid) | Vc_ComFB-His_Fw:<br>CTAGAAATAATTTTGTTTAACTTTAAGAAGGAGATATACCATGCAATTGAGCGAAGATT | This study |
| | Vc_ComFB-His_Rev:<br>CAGCCGGATCTCAGTGGTGGTGGTGGTGGTGGTGGTGCGACTTATGGTGCTGAACTTGGTTG | |
| $His_8$-tagged *Tb*ComFB (pET28a_*Tb*ComFB-$His_8$ plasmid) | Gene Block encoding *Thermoanaerobacter brockii* ComFB<br><br>CTAGAAATAATTTTGTTTAACTTTAAGAAGGAGATATACCATGCAACTTAAAAATTACATGGAAGACGCTGTTGAG<br>CAGATGATGGACAGGGTTTTAAAAGACCTGGACGTTTGTAAGTGTGACAGGTGTAGGATGGACATAAAAGCTTTA<br>GCCCTTAACAATTTGCCTCCGAAATATGTTGTATCAGAAGAAGGAGAACTGTATGTAAAGACAAACGAGTTAGTCA<br>GGCAATTTGAGGTAGACATAATAAAAGCCATAACAATGGCTGCTATTAAAGTGAATAACAATAAGCGCCATTCGC<br>ATCATCACCACCACCACCACCACTGAGATCCGGCTGCTAACAAAGCCCG | This study |
| $His_8$-tagged *Td*ComFB (pET28a_*Td*ComFB-$His_8$ plasmid) | Gene Block encoding *Treponema denticola* ComFB<br><br>CTAGAAATAATTTTGTTTAACTTTAAGAAGGAGATATACCATGATCATTCATAACGTTATGGAAGACTTGGTATATA<br>CCGAAGTCAACAAACTATTTGATGAAGCCGAAGAAAAAAAAGAAAGCTGGCTTACATGCAGTTGTATGCAATGCA<br>GAGTTGACACCATGTGCTATGTACTCAACCGGGTTAAACCCCGCTATATAAAATCGGGGAGAGGCCTCGCTCACTT<br>TTTAAAATTCGAAAAAACGGAAAAAGTACAGATTATGGCCGATATTACAAGTTTGGTAATTGAAGGTATGCACAG<br>GGTTCTTTCTACAAAAAGGCCTCATGACCATGAACCCATATTCGAAACCGAAAACACACCCGTTTTTAATTTTCCTG<br>CCATAACAGGCAATATTTTAAACGGAAGCAATTTTAGACCGATGGAAGATGCTACGATAAGCCTTAAAATGAACG<br>GAGAGCTGGTATCCCAGATGAGCATTCTTTGGGATAATCCTTATACAATATCGGAAAAGACACCCGGAGCTTATAC<br>CTTTTGTCCCAAAGCAATTCCTGCAAAAAATGCAGGAGATAAGGAAAAGTTTATTTTTGTCCTTAAAGCTGAAAAA<br>GAAGGCTTTGATCCGACTAACTTTTCATTCGATATTGAATTAACGGCAGAAGACATGGTAAAATCTCCTTTGGACTC<br>TTCAAATTTTTATCAAATCAAAAATTTGTTCTTATGCGAACACAGTGAGGAAGAATCGCATCATCACCACCACCACC<br>ACCACTGAGATCCGGCTGCTAACAAAGCCG | This study |
| pET28a_sequancing primers | 5'-pET: CCGCGAAATTAATACGACTCAC | |
| | 3'-pET2: CCTCAAGACCCGTTTAGAGG | |
| $P_c$-FB forward | 5′-GCCATG**AAGCTT**AAAGGAGGAAACAATCATGCTTGTCAATTCAAAAGA | This study |
| $P_c$-FB-reverse | 5′-ATCGCC**GGATCC**GCG AATCACATAATAAACAGATCA | This study |
